# Genomic insights into post-domestication expansion and selection of body size in ponies

**DOI:** 10.1101/2023.08.25.554910

**Authors:** Xingzheng Li, Zihao Wang, Min Zhu, Binhu Wang, Shaohua Teng, Jing Yan, Pengxiang Yuan, Shuwei Cao, Xiaolu Qu, Zhen Wang, Panir Choudhury, Xintong Yang, Qi Bao, Sang He, Lei Liu, Pengju Zhao, Jicai Jiang, Hai Xiang, Lingzhao Fang, Zhonglin Tang, Yuying Liao, Guoqiang Yi

## Abstract

Horses domestication revolutionized human civilization by changing transportation, farming, and warfare patterns. Despite extensive studies on modern domestic horse origins, the intricate demographic history and genetic signatures of pony size demand further exploration. Here, we present a high-quality genome of the Chinese Debao pony and extensively analyzed 385 individuals from 49 horse breeds. We reveal the conservation of ancient components in East Asian horses and close relationships between Asian horses and specific European pony lineages. Genetic analysis uncovers Asian paternal origin for European pony breeds, and these pony-sized horses share a close genetic affinity due to the presence of a potential ancestral ghost pony population. Additionally, we identify promising cis-regulatory elements influencing horse withers height by regulating genes like *RFLNA* and *FOXO1*. Overall, our study provides insightful perspectives into the development history and genetic determinants underlying body size in ponies and offers broader implications for horse population management and improvement.

**Teaser:** Decoding pony genetics: exploring origins and size determinants sheds light on their historical and biological impacts.

## Introduction

The domestication of animals played a pivotal role in shaping human civilization. Horses were first domesticated approximately 5,500 years ago on the Eurasian steppes(*1*). Their use in transportation, agriculture work, and warfare facilitated the expansion of human territories and the establishment of long-distance trade routes(*2–4*). Among the diverse group of horse breeds, ponies are a type of small horses (*Equus caballus*), which are generally defined as individuals that stand less than 14.2 hands (58 inches, 147 cm) at the withers. They have been selectively bred for their adaptability to diverse local environments(*5*), ranging from the islands of Northern Europe to the mountain regions of Southern China. Due to their hardness, intelligence, and friendly nature, ponies served various purposes including transportation, work, and companionship(*6*). In recent years, they have also gained popularity as mounts for children. Despite the important roles that ponies have played in human history, our knowledge of the genetic factors contributing to their unique characteristics remains limited. Therefore, investigating the genetic architecture and diversity of pony breeds could help to provide novel insights into the domestication history of horses and trace human activity and civilization.

As one of the most important local horse breeds from Guangxi province in China, the Debao pony reaches a height of around 9.3 hands (39 inches, 100 cm) at the withers and is believed to be a descendant of ancient breeds that existed 2,000 years ago. While previous study has provided insight into the origins and spread of domestic horses from the Western Eurasian steppes(*1*), the development history of horses after domestication, especially that of ponies in early times and their relationships, remains unclear. In particular, the genealogy of pony clusters from Southern China among horse breeds remains uncertain due to the lack of a comprehensive dataset. Additionally, many previous studies have identified a series of candidate genes (*TBX3*, *NELL1*, *ZFAT*, *LASP1*, *LCORL/NCAPG*, *ADAMTS17*, *GH1*, *OSTN*, and *HMGA2*) associated with body size in horses(*7–11*). Still, these findings are limited by a small subset of breeds or geographic regions. Therefore, further integrative analysis, including Eurasian pony breeds, would generate new avenues for understanding the common genetic basis that shapes the characteristic and diversity of ponies.

To enhance our understanding of the complex demographic history and genetic signatures of body size in ponies, we first assembled a high-quality Debao pony genome by combining PacBio CLR, Illumina sequencing, and Hi-C technologies. Subsequently, we gained insight into the development and breeding history of ponies by performing a comprehensive whole-genome analysis based on 385 domesticated horse individuals from 49 breeds in the world. Through integrating genetic, 3D genome, and histone mark ChIP-Seq datasets, we uncovered novel genetic factors influencing the withers height of horses. This study provides new perspectives on the horse development history post domestication and has important implications for the management and improvement of modern horse populations.

## Results

### Genome assembly and annotation of Debao pony genome

We assembled a chromosome-level genome of Debao pony by integrating PacBio CLR (296 Gb, 121×), Illumina PE (333 Gb, 136×), and Hi-C (119 Gb) sequencing technologies (Fig. 1A and fig. S1). The estimated genome size was 2.44 Gb with a heterozygosity of 0.29% (fig. S2 and table S1). By conducting self-correction, trimming, and assembly of PacBio long reads with Canu (fig. S3), we initially obtained 583 contigs with an N50 of 20.48 Mb after purging haplotigs (table S2). The contigs were then anchored, ordered, and orientated into a chromosome-level assembly using the 3D-DNA pipeline. The resulting assembly underwent manual curation and polish (fig. S4). Each step went through rigorous quality checks to ensure accuracy (fig. S5). Hi-C signal heatmap demonstrated clear genomic interaction, suggesting high quality and accuracy (Fig. 1B and fig. S6). The final genome assembly spanned 2.44 Gb, consisting of 31 autosomes, a pair of X/Y sex chromosomes, a mitochondrial genome, and 210 unassigned contigs, with a contig N50 of 34.25 Mb and a scaffold N50 of 88.98 Mb (Fig. 1C and table S2). Notably, the Debao assembly rectified un-collapsed artificial duplications present in EquCab3.0 (Fig. 1D), exhibiting high contiguity and completeness among released *E*. *caballus* genomes (Fig. 1, C and E, fig. S7 and table S2).

**Fig. 1.**
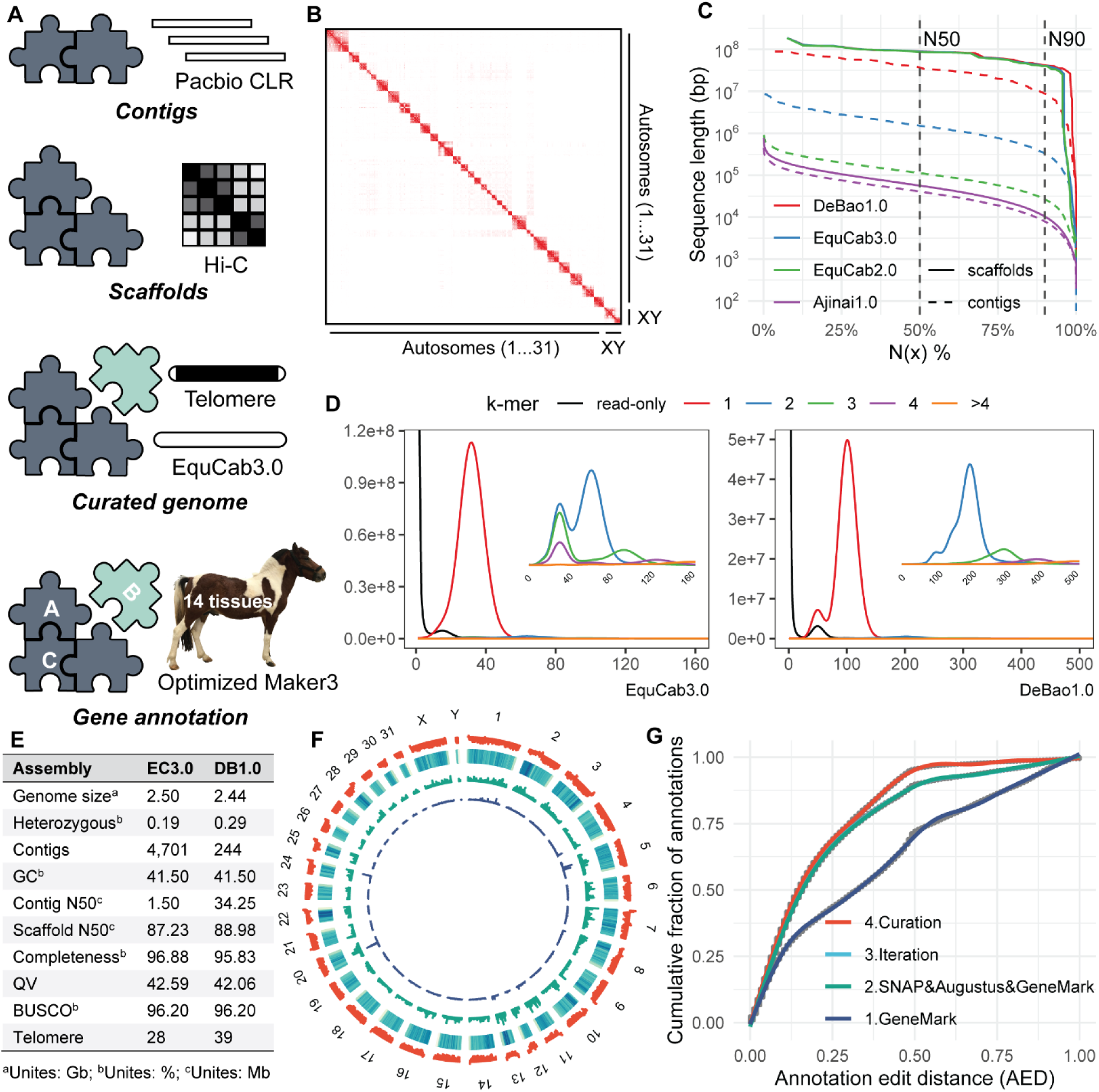
Genome assembly and annotation of the Debao pony. (**A**) Schematic diagram illustrating the pipeline for genome assembly and annotation. (**B**) Hi-C chromatin interactions of the assembled pony genome. (**C**) Comparison of assembly contiguity between current released horse genomes and DeBao1.0 assembly. (**D**) Copy number spectrum (spectra-cn) of k-mers for EquCab3.0 and DeBao1.0. Spectra-cn of k-mers larger than 1 (2, 3, 4, >4) are enlarged within each plot. (**E**) Assembly statistics of EquCab3.0 and DeBao1.0. The contig N50 was calculated by breaking the genome into contigs at the gap region. (**F**) Landscape of the assembled pony genome, showing chromosome names, GC contents, repeat density, gene density, and ncRNA density in different tracks from outer to inner. (**G**) Annotation edit distance (AED) using iterative Maker3 annotation to assess annotation accuracy.

Repetitive sequences accounted for 41.26% of the Debao genome using Dfam, Repbase, and Repeatmodeler (table S3). We predicted a total of 21,038 high-confidence protein-coding genes based on an optimized MAKER annotation pipeline that combined *ab initio* gene prediction, protein-based homology searches, RNA-Seq, and Iso-Seq (Fig. 1, F and G, and fig. S8). The gene models were further refined using Apollo. Notably, utilizing the assemblies of Debao pony and thoroughbred as representatives of *E. caballus*, we revealed a significant gene loss associated with olfactory receptor activity, a phenomenon observed across diverse horse breeds(*12*). This finding suggests that this gene family may undergo relaxed selective constraints during horse domestication (fig. S9). The high quality of the Debao pony genome establishes it as a valuable resource for genetic research.

### Genome-wide variation and population structure of *E*. *caballus*

To gain a comprehensive understanding of the genetic architectures of ponies across horse populations, we re-sequenced 10 Debao ponies, 10 Baise horses, and six Warmbloods at an average depth of approximately 10.87×. Then, we collected 384 previously sequenced genomes to capture more genetic diversity in horses. After excluding potential crossbred samples, we obtained a panel of 385 samples representing 49 horse breeds worldwide (Fig. 2A and tables S4 and S5). Using the Genome Analysis Toolkit (GATK), we identified 35,926,564 high-quality SNPs throughout the genome (fig. S10 and tables S6 and S7), with 60.60% located within intergenic regions and 27.36% within introns (table S8).

**Fig. 2.**
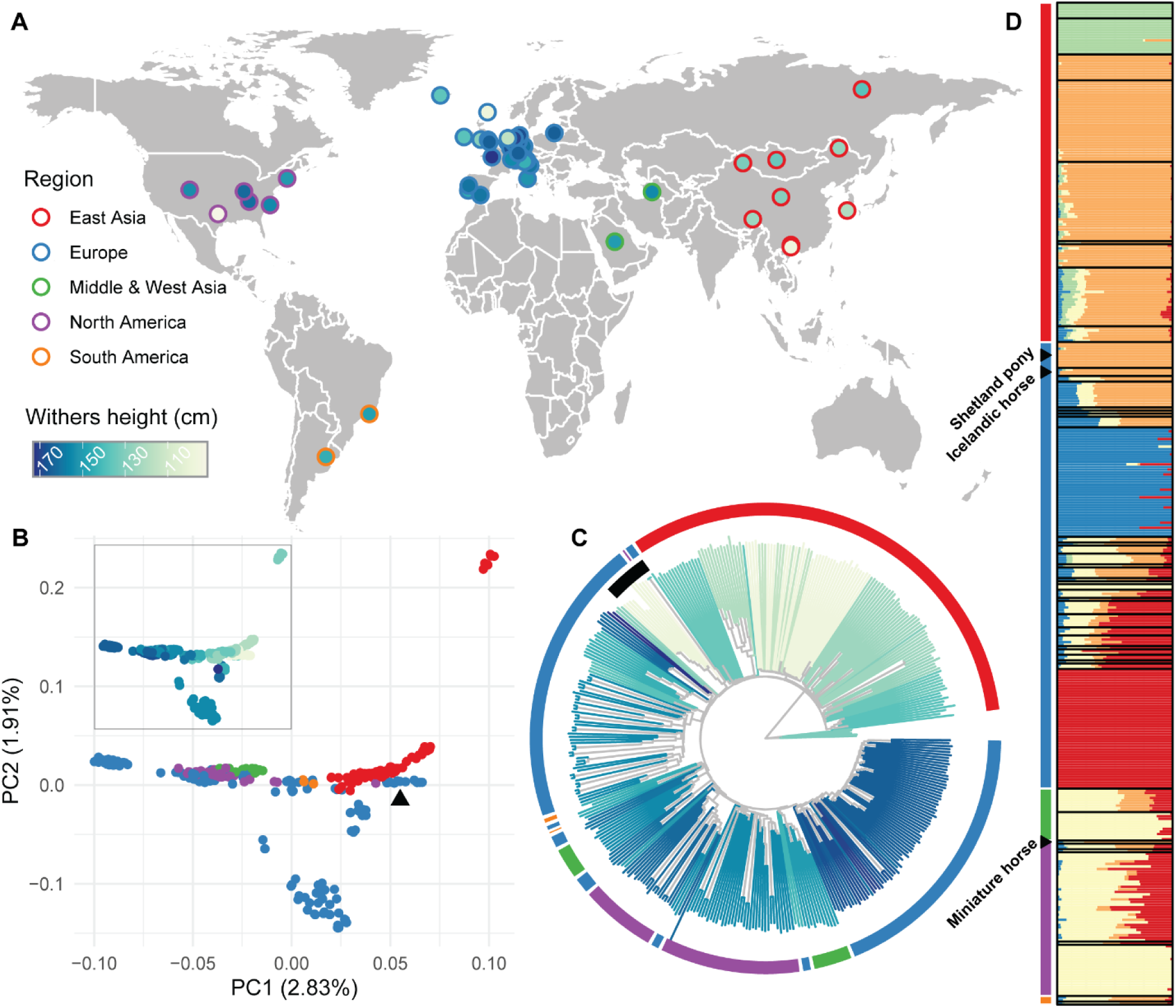
Geographic distribution and population structure of *E. caballus*. (**A**) Geographic origin of 49 horse breeds, with colors representing regions and withers height consistent throughout the study. (**B**) Principal component analysis (PCA) plot showing the genetic relationships among horse populations. The black triangle indicates the European pony lineage, which includes the Shetland pony, Icelandic horse, and Miniature horse. The colors of the data point in the main plot and the thumbnail (top left) represent regions and withers height, respectively. (**C**) Neighbor-joining (NJ) tree rooted on Przewalski’s horses depicting the genetic clustering of horse populations. The black inner section ring highlights the European pony lineage of Shetland pony, Icelandic horse, and Miniature horse. Colors of the outer section ring indicate regions, while branch colors represent withers height. (**D**) Admixture analysis of horse populations (*K* = 5), revealing genetic contributions from different ancestral sources. Breeds are ordered by regions and are represented by strip colors.

All eligible SNPs were employed to investigate population structure and diversity within the horse population. Principal component analysis (PCA) exhibited geographic clustering and a clear spectrum reflecting the gradient change of withers height (Fig. 2B and table S9). Notably, horses in East Asia were generally pony-sized, and ponies from other regions, especially those belonging to the lineage containing Shetland pony, Icelandic horse, and Miniature horse, exhibited relatively closer relationship with East Asian horses. Furthermore, the neighbor-joining (NJ) phylogenetic tree displayed similar classification patterns, albeit with relatively scattered geographic clustering due to complex breeding history (Fig. 2C). The pony lineage of Shetland pony, Icelandic horse, and Miniature horse showed phylogenetic proximity to East Asian horses. These findings were also strongly supported by the admixture results. With an optimized *K* value of five, the European pony lineage predominantly shared more ancestry components with Asian horses rather than European horses (Fig. 2D and fig. S11). With higher *K* values, this lineage segregated from Asian horses and formed a distinct group (fig. S12 and table S10). Our analysis reveals the genome-wide variation and population structure of *E. caballus* and provides novel insights into the close relationship between the specific European pony lineage and the East Asian horse.

### Y chromosome haplotypes (HTs) and phylogenetic analysis

To investigate the paternal genetic history of pony populations, we analyzed Y chromosome haplotypes and phylogeny using 177 male individuals in our horse dataset (fig. S13 and table S11). In total, we identified 59 diverse haplotypes, and Przewalski’s horse harbored a unique haplotype distinct from all domestic horses (Fig. 3A and table S12). The haplotype network comprised three major haplogroups (II, V, and VI), with Asia horses like Debao pony, Mongolian horse, and Tibetan pony displaying the highest haplotypes diversity, which surrounded the ancient HT II (Fig. 3B). Notably, we found that the paternal lines of Shetland pony and Icelandic horse originated from HT II and its closely related haplotypes, respectively. Furthermore, the phylogenetic analysis revealed that the paternal lineages of the Shetland pony and Icelandic horse were primarily descended from horses in Asia, suggesting an orient paternal origin (Fig. 3C). Overall, our analysis by tracing paternal lineage ancestry provides further evidence for the genetic diversity and evolutionary history of ponies.

**Fig. 3.**
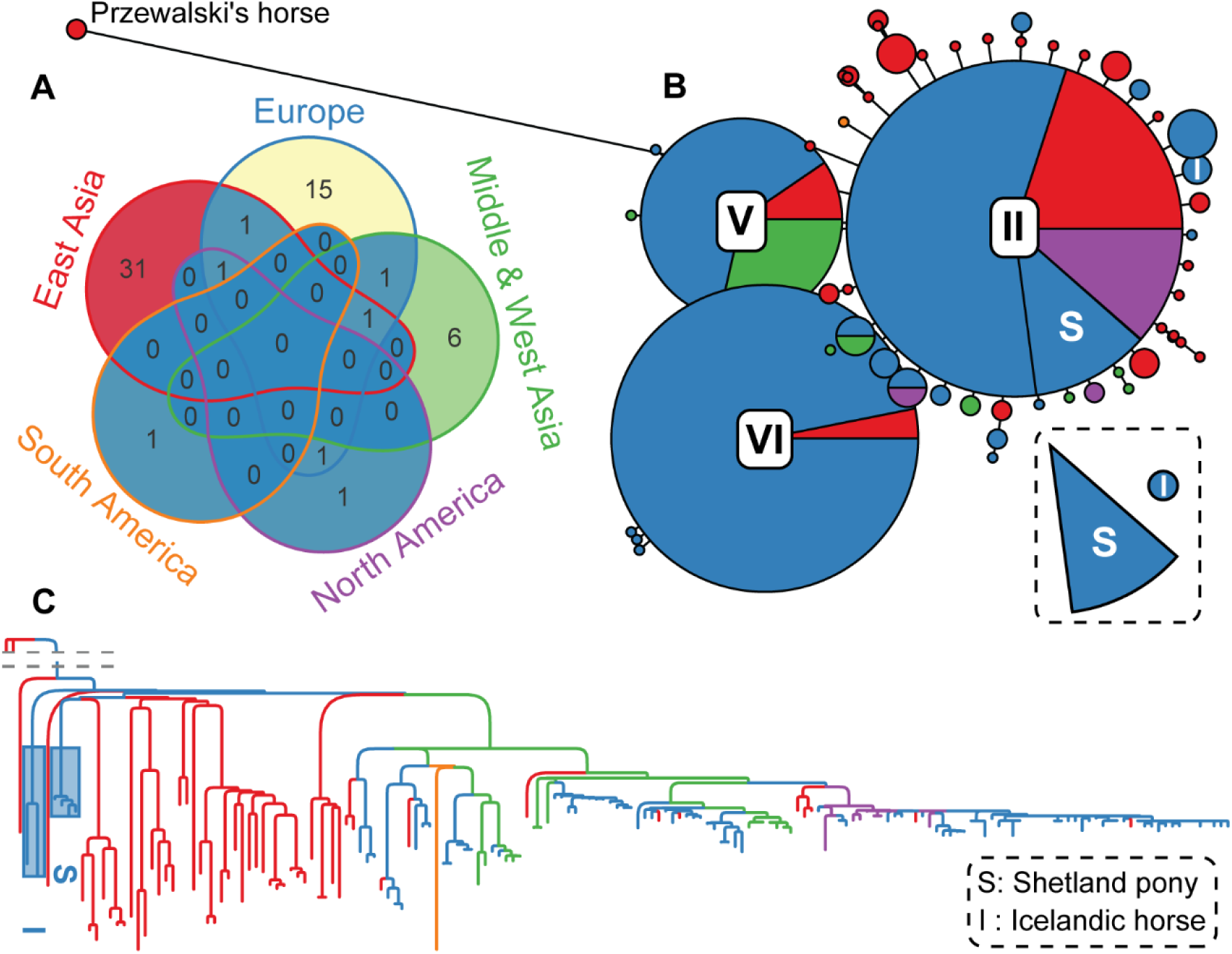
Characterization of the Y chromosome in stallions. (**A**) Y chromosome haplotypes in a total of 177 horses across 38 breeds. (**B**) Haplotype networks constructed based on Y chromosome segregating sites. Circle size represents the frequency of each haplotype, while the length of lines corresponds to the number of mutations. (**C**) Hierarchical clustering based on variations observed in the Y chromosome. The letters “S” and “I” represent the Shetland pony and the Icelandic horse, respectively.

### Phylogenetic reconstruction, genetic affinities, and demographic inferences of ponies

To further evaluate the amount of shared genetic drift between any two horse populations, we carried out phylogenetic reconstruction and outgroup *f3*-statistics analysis. Heatmaps of outgroup *f3*-statistics showed early breeding of Asian horses, followed by European ponies, which aligned with our population structure results (Fig. 4A and figs. S14 and S15). The rapid linkage disequilibrium (LD) decay of ponies also suggested a more primitive state (Fig. 4B). To capture global relationships as well as developmental trajectories of different breeds, we calculated the likelihood of transition between states by employing the potential of heat diffusion for affinity-based transition embedding (PHATE). The PHATE-embedded data generated nine major linear trajectories representing divergent genetic states, which extended from the central portion (Fig. 4C and table S13). Notably, the Shetland pony formed one of the major trajectories. Moreover, the two Debao ponies were positioned at the basal portion, reflecting transitional genetic states between the Shetland pony and their wild ancestry. Investigation using TreeMix inferred gene flow points from the Yakutian horse to both the Shetland pony and the Debao pony (Fig. 4D and table S14). To further classify gene flows underlying pony domestication and breeding history, we calculated *f4*-statistics for every possible combination across all sequenced breeds and constructed a pony relationship network based on key *f4*-statistic values (fig. S16). The network indicates that gene flows and connections between western and eastern ponies were mainly mediated by the Yakutian horse (Fig. 4E and fig. S17). Based on geographic distance and prior knowledge of Yakutian horse origin, we speculated that the majority of modern ponies descended from an ancient unidentified pony lineage rooted in DOM2, which was closely related to the Yakutian horse. Demographic inference using SMC++ displayed an early bottleneck in eastern pony populations, followed by western pony populations with lower effective population sizes (*N_e_*), supporting early breeding of eastern ponies and subsequent westward dispersal of a small fraction of the pony population (Fig. 4F). Altogether, we hypothesized that the propagation of contemporary ponies primarily originated from a shared ancestral pony population within DOM2 in the Eurasian steppe (Fig. 4G).

**Fig. 4.**
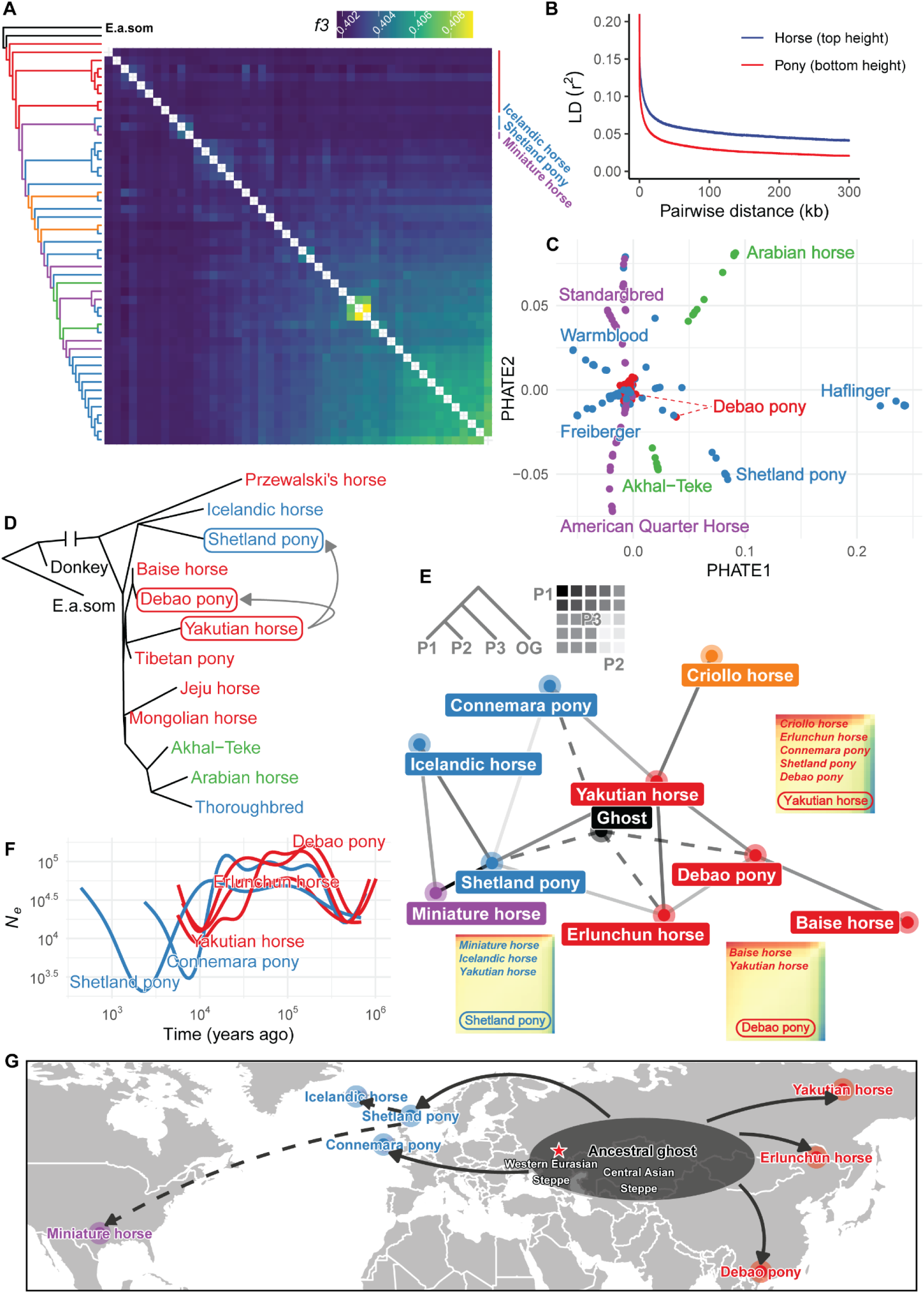
Genetic affinities of ponies in the horse population. (**A**) TreeMix Phylogenetic relationships and outgroup *f3*-statistics (*Equus caballus*, *Equus caballus*, *E.a.som*) of the horse population. (**B**) Linkage disequilibrium (LD) decay for the top-sized horses and bottom-sized ponies. (**C**) PHATE plot depicting distinct horse breeds. (**D**) Phylogeny and inferred mixture events by TreeMix. (**E**) Pony-related network based on *f4*-statistics. Line colors ranging from light to dark indicate genetic relationships from distant to close. The top left thumbnail illustrates the basic unit of *f4*-statistics. Examples of P3, such as the Yakutian horse, Shetland pony, and Debao pony, are displayed surrounding the main network. The most genetically closely related breeds for each P3 are listed starting from the top left. (**F**) Demographic trajectories of pony populations inferred by SMC++. (**G**) Modeled dispersal of ponies based on sequencing data analysis and historical knowledge. The red star represents the inferred horse domestication center.

### Novel selection signatures associated with horse withers height

To decipher the genetic basis of body size variation in horses, we performed a comprehensive genome-wide selection analysis by focusing on three pony breeds and eleven horse breeds with divergent withers heights (Fig. 5A). By integrating multiple selection metrics including fixation index (*F*_ST_), nucleotide diversity (π) ratio, and cross-population extended haplotype homozygosity (XP-EHH), we identified numerous candidate genomic regions and associated genes (Figs. 5, B and C, and tables S15 to S17). Notably, the genes covered by the selective regions identified by these three methods exhibited significant enrichment in the Dorso-ventral axis formation KEGG pathway, highlighting the role of early developmental processes in shaping body size differences (Fig. 5, C and D, and table S18). Within these candidate regions, we discovered several genes previously implicated in horse body size, such as *HMGA2* and *TBX3*. Additionally, our analysis revealed several novel candidate genes, including *ACSF3*/*CENPBD1* (genomic region), *RFLNA*, *KIF2B*, *FOXO1*, and *ABT1*, which are potentially involved in regulating horse withers height (Fig. 5B).

**Fig. 5.**
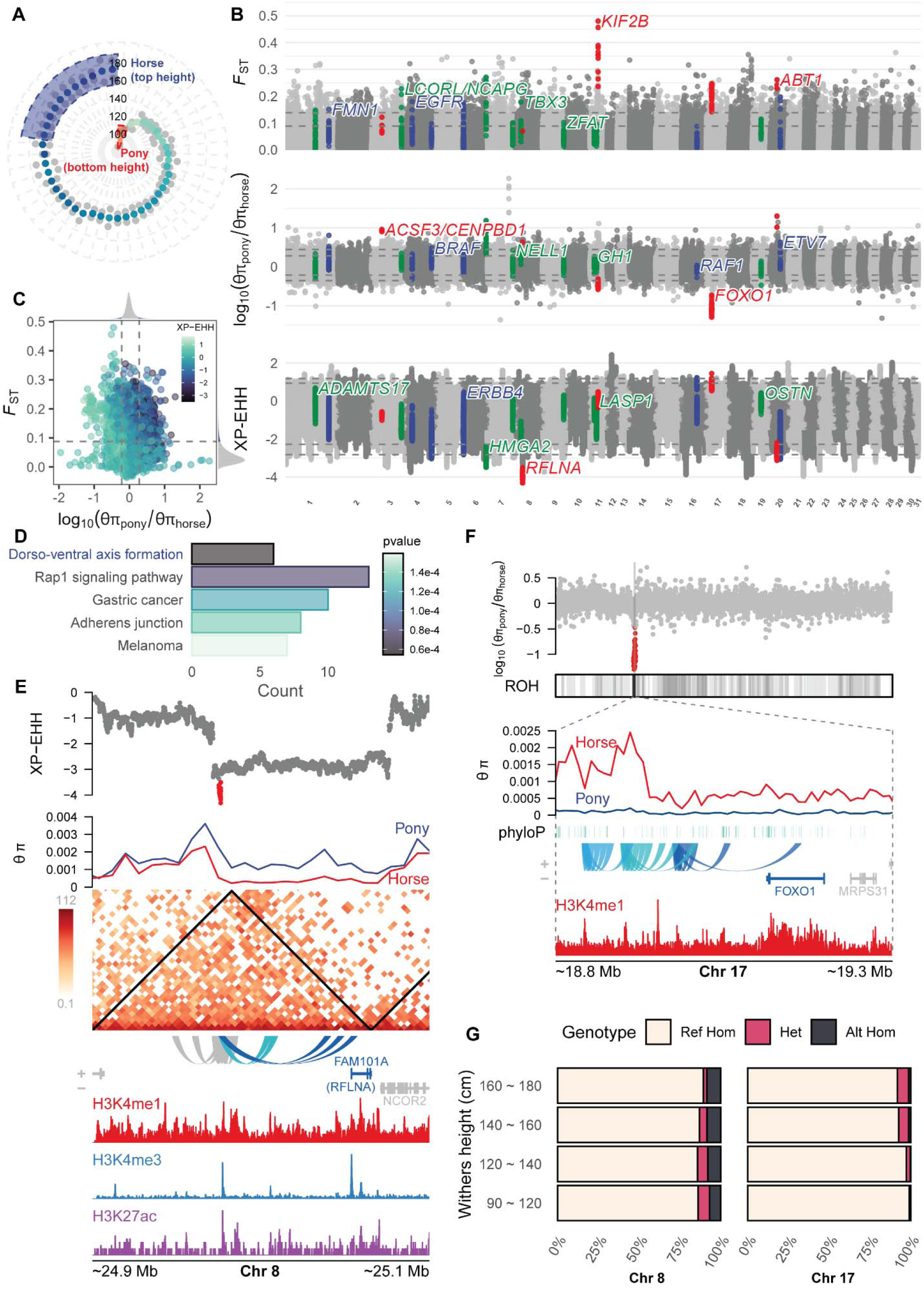
Selective regions associated with horse withers height. (**A**) Breeds representing large-size horses and small-size ponies. Horses (top height): Shire horse, Percheron, Oldenburger, Holsteiner, Württemberger, Hanoverian horse, Dutch Warmblood, Westphalian horse, Bavarian Warmblood, Warmblood, and Trakehner. Ponies (bottom height): Shetland pony, Debao pony, and Miniature horse. (**B**) Genome-wide selective signals associated with horse body size based on *F*_ST_, π ratio, and XP-EHH (pony vs. horse). The top 1% and 5% quantiles are indicated by grey dashed lines. Red points represent candidate regions, blue points represent gene regions enriched in the Dorso-ventral axis formation KEGG pathway, and green points represent previously reported gene regions associated with horse body size. (**C**) Overview of selective signals incorporating *F*_ST_, π ratio, and XP-EHH. The top 5% quantiles are indicated by grey dashed lines. (**D**) Top enriched KEGG pathways. (**E**) Candidate selective region on chromosome 8. From top to bottom are XP-EHH, θπ, Hi-C matrix, genomic interactions, gene models, and histone ChIP-Seq (diaphysis of metacarpal bone) signals. TADs are indicated with black triangles in the Hi-C matrix. (**F**) Candidate selective region on chromosome 17. From top to bottom are π ratio (θπ_pnoy_/θπ_horse_), ROH_pony_, θπ, phyloP, genomic interactions, gene models, and histone ChIP-Seq (diaphysis of metacarpal bone) signals. (**G**) Genotype frequency of candidate regions at different withers heights located on chromosomes 8 and 17, respectively.

One of the identified selective regions corresponds to a regulatory element located upstream of the *RFLNA* gene, which is known to be involved in the regulation of bone mineralization, bone maturation, and chondrocyte development, spanning approximately 1.3 kb on chromosome 8 (Fig. 5E). This region exhibited the lowest XP-EHH values (pony vs. horse) across the entire genome, indicating positive selection for large body size. Moreover, the horse populations displayed reduced nucleotide diversity in this genomic region, further supporting its role in selection for body size. By integrating three histone marks ChIP-seq data from the FAANG database, we revealed the potential enhancer function for the regulatory element of *RFLNA*, which is supported by the topologically associating domain (TAD) inferred from Hi-C data.

Another identified selective region harbored multiple enhancers located downstream of the *FOXO1* gene, which is known regulate osteoblast numbers, bone mass, and chondrogenic commitment of skeletal progenitor cells, spanning from 18.75 to 19.08 Mb on chromosome 17 (Fig. 5F). The presence of low nucleotide diversity and extended runs of homozygosity (ROH) in the pony population suggests the conservation of this region across most pony breeds, emphasizing its significance in determining small body size (fig. S18). These closely positioned enhancers, along with the proximity to the *FOXO1* gene, suggest their potential synergistic regulation of *FOXO1*. The consecutive blocks exhibiting high phyloP scores further support its pivotal role.

In addition to the aforementioned instances, several other selective signals were associated with body size (figs. S19 to S21). Given that body size is a polygenetic trait, we showed the difference in the allele frequency spectrum at candidate loci regarding four groups of body size. The presence of a gradient trend in allele frequencies across these selective regions, in concordance with withers height, further emphasizes their association with horse body size (Fig. 5G and fig. S22). Collectively, these findings underscore the significance of the regulatory elements flanking the candidate genes and their potential contribution to the variation in horse body size.

## Discussion

The origin and domestication of modern horses have long fascinated researchers and garnered significant interest(*13–15*). The Western Eurasian steppes, particularly the lower Volga-Don region, have been identified as the probable domestication center of modern domestic horses known as DOM2, potentially dating back to around 3000 BC(*1*). This marked the beginning of their global expansion, resulting in the emergence of diverse horse breeds with a wide range of body sizes, varying from less than 1 meter to nearly 1.8 meters at the withers. Among these horse breeds, ponies have gained increasing popularity worldwide due to their small size, versatility, appeal to various age groups, accessibility, and therapeutic benefits(*6*). However, their genetic relationships and breeding history have remained largely unknown. To gain valuable insights into the development history and specific characteristics of ponies, we assembled the pony genome and conducted subsequent analyses using global sequencing data.

In contrast to using short-read data in EquCab3.0(*16*), we directly employed Pacbio long-read data to construct the genome backbone. This approach yielded remarkable improvements in contiguity, with an impressive 20-fold increase in the contig N50 value. Moreover, we achieved a base accuracy comparable to that of EquCab3.0, which was established from the foundation of Sanger sequence data(*16*). Importantly, the current assembly also addressed the issue of false repetitive sequences that were overrepresented in EquCab3.0, potentially causing overestimated gene family expansions(*17*). The resulting high-quality pony genome enhances our understanding of horse genetic diversity and provides the scientific community with a valuable resource for horse genetics and genomics. Moving forward, the pony genome along with other horse assemblies can be employed for large-scale pan-genome analysis, which would not only enrich the genome-wide genetic variations but also shed light on the biological adaptability across diverse horse breeds(*18*).

Population structure analysis revealed that the western pony lineage, consisting of the Shetland pony, Icelandic horse, and Miniature horse, exhibited a closer genetic relationship to oriental horses. This suggests potential historical connections and gene flow between these pony breeds and oriental horse populations. Historically, horses have been transported and traded through various means, including invasion, immigration, or commerce(*19–21*). Previous evidence suggests that early breeders likely selected stallions for breeding based on visible phenotypic traits(*22*). Therefore, we delved deeper into the genetic history by investigating the Y chromosome lineages, providing compelling evidence supporting an oriental paternal origin for this European pony lineage. East Asia, particularly the southwestern region of China, has preserved a rich diversity of Y chromosomal haplotypes and ancient paternal horse lineages(*23*). In contrast, Europe has experienced a decline in genetic diversity among ancient domestic stallions over the past 4,000 years. Notably, Y-HT-1 (haplotypes exhibited by present-day horses) has replaced most European haplotypes, except for Yakutian horses, which suggests a higher diversity of patrilineages originating from further East(*24–26*). However, limited ancient materials have been preserved or found to reveal the landscape of ancient horse Y chromosomes in East Asia. In our research, we further elucidated the ancient component of East Asian horses and demonstrated a genetic connection between European and Asian ponies through Yakutian horses. Contemporary Yakutian horses were most likely introduced following the migration of the Yakut people from the Altai-Sayan and/or Baïkal area between the 13^th^ and 15^th^ centuries or even earlier(*27*). The fast adaptation and geographical isolation of Yakutian horses helped maintain abundant ancestral components in their genome. Previous studies estimated that the Shetland pony and Icelandic horse originated over 1,000 years ago, accompanying the arrival of Norse settlers during the Viking Age(*28*).

The Debao pony is believed to be a descendant of the ancient Guoxia breed, which dates back thousands of years. However, the exact origins of their ancestors remain elusive and cannot be traced with certainty. War horses, known for their large size and high speed, facilitated the spread of humans and horse breeds through warfare(*29*). However, pony-sized horses do not possess the same physical attributes or speed advantage. Therefore, the breeding history and spreading route of ponies may be more complex and obscured by the intricate relationships among horses. Based on our comprehensive analyses, we speculated that they originated from the ancestral ghost pony population within DOM2, likely centered around the Central Asian Steppe. The absence of the ancestral ghost population and the close affinity of Yakutian horses with it results in the current genetic relationship network. Interestingly, these pony-sized breeds are all located on islands or in mountainous regions, where isolated geography has hindered gene exchange to some extent, leading to their basal localization in the phylogeny. Given that previous studies have predominantly focused on European horses(*24*), these findings extend our understanding of the ancient component and genetic diversity of East Asia horses. To fully understand the breeding history of ponies and other horse breeds post-domestication, additional ancient genomes covering a wider geographical range, especially in East Asia and local rural areas, will be crucial.

While previous work identified candidate genes associated with body size, most of these studies focused on specific breeds/regions or relied on low-density Chip datasets(*8, 10*). By integrating a large-scale resequencing dataset encompassing horses with a wide range of heights, we were able to uncover previously unknown genetic information related to body size. For instance, frameshift variants in *RFLNA* have been linked to spondylocarpotarsal synostosis syndrome, a skeletal disorder characterized by short stature and carpal/tarsal synostosis(*30*). The involvement of *FOXO1* in embryonic development, bone growth and remodeling, and cartilage repair processes underscores its essential function in skeletal development(*31*). Additionally, the *ACSF3/CENPBD1* genomic region showed enrichment of genes, including *ACSF3*, *ANKRD11*(*32*), *CDK10*, *TCF25*, *DEF8*, and *CENPBD1*, involved in body development (fig. S19). Another signal was identified near *KIF2B* gene, which belongs to the KIF family involved in the development of the nervous system and early embryo (fig. S20). Furthermore, a selective signal encompassed regulatory elements of the *ABT1* gene, which plays a critical role in growth and height trait (fig. S21)(*33*). One interesting finding is that most candidate regions were located within cis-regulatory elements, underscoring the significant role of transcriptional evolution in driving rapid adaptation and the breeding process. These newly discovered cis-regulatory elements not only provide insights into the function of noncoding regions but also deepen our understanding of the regulation of horse height. Further functional characterization and validation of these genes will contribute to our understanding of the molecular mechanisms driving withers height variation in horses.

In conclusion, our study assembled and annotated a high-quality pony genome, providing a valuable resource for future horse research endeavors. Through comprehensive genetic analysis, we gained profound insights into the intricate genetic relationships among various pony breeds. Additionally, our investigation into the horse withers height led to the discovery of novel cis-regulatory elements involved in this trait. These findings enhance our understanding of pony genetics and provide a foundation for future studies on breed characteristics and trait selection.

### Materials and Methods Ethics statement

Our study adhered to the ethical guidelines outlined in the Guide for the Care and Use of Experimental Animals, as established by the Ministry of Agriculture and Rural Affairs (Beijing, China). The protocols were reviewed and approved by the Institutional Animal Care and Use Committee of both the Chinese Academy of Agricultural Sciences and the Guangxi Veterinary Research Institute. To minimize any potential suffering, horses were humanely euthanized as required prior to tissue sampling.

### Genome assembly

A male Debao pony from Debao County, Baise, Guangxi Province, China, was selected for genome assembly. Genomic DNA was extracted from its blood using the DNeasy Blood & Tissue Kit (Qiagen, Beijing, China), and DNA quality was assessed by agarose gel electrophoresis. For long-read sequencing, single molecular real-time (SMRT) PacBio sequencing libraries were prepared following standard Pacific Biosciences protocols and sequenced on the PacBio Sequel platform. Hi-C libraries were constructed using Phase Genomics Proximo Hi-C Animal Kit (Phase Genomics, Washington, USA). Following the manufacturer’s instructions, chromatin was digested with the DpnII restriction enzyme, and the Hi-C libraries were sequenced on an Illumina NovaSeq 6000 platform. Additionally, short-read paired-end (PE) sequencing libraries were prepared and sequenced on an Illumina NovaSeq 6000 platform.

The genome assembly followed the Vertebrate Genomes Project (VGP) assembly pipeline(*34*) with specific modifications (fig. S1). The genome size of the Debao pony was estimated by analyzing the frequency distributions of 17, 19, 21, 23, 25, 27, 29, and 31 mers from Illumina PE reads using jellyfish (v2.2.10)(*35*) and calculated using GenomeScope (v2.0)(*36*). The PacBio reads were independently assembled into contigs using three different assemblers: FALCON (v0.3.0)(*37*), wtdbg2 (v2.5)(*38*), and Canu (v2.1.1)(*39*). Redundant sequences from the Canu assembly were removed using purge_dups (v1.2.5)(*40*). The contigs generated by Canu were then scaffolded based on continuity and completeness (fig. S3 and table S2), which used the 3D-DNA pipeline(*41*) with Hi-C data preprocessed by fastp (v0.20.1)(*42*). The genome assembly was manually curated using PacBio long reads, telomere locations, Hi-C signals, and synteny information with the EquCab3.0 reference genome, aided by Juicebox Assembly Tools (v1.11.08)(*43*). The curated genome, along with the mitochondrial genome generated by GetOrganelle (v1.7.5)(*44*), was polished through two rounds of PacBio long reads using the Arrow algorithm and two rounds of Illumina short reads using Pilon (v1.24)(*45*). Chromosome numbers were assigned based on the alignment with EquCab3.0 (fig. S5). Throughout the genome profiling stage, the assembly quality was evaluated using BUSCO (v5.2.2, mammalia_odb10)(*46*), Merqury (v1.3)(*47*), MUMmer (v4.0.0rc1)(*48*), and QUAST (v5.0.2)(*49*).

### Genome annotation

Repeat sequences were identified using a combination of homology-based and *de novo* approaches. The homology-based method employed the RMBlast (v2.9.0, http://www.repeatmasker.org/rmblast/) search engine with the transposable element (TE) database from Dfam (v3.2, https://dfam.org) and Repbase (v20181026)(*50*). *De novo* prediction was performed using RepeatModeler (v2.0.1)(*51*), which integrated TRF (v4.09)(*52*), RECON (v1.08)(*53*), RepeatScout (v1.0.6)(*54*), and LTR_Retriever (v2.9.0)(*55*). The merged TE library was utilized for repeat sequence identification using RepeatMasker (v4.1.1, http://www.repeatmasker.org).

For genome annotation, an optimized iterative approach using MAKER3 (v3.01.03) was employed (fig. S8)(*56, 57*). The repeat library generated above was used to mask the genome. Evidence for gene annotation included protein sequences of six species (*Bos taurus*, *Equus caballus*, *Homo sapiens*, *Mus musculus*, *Ovis aries*, and *Sus scrofa*) retrieved from UniProt (https://www.uniprot.org/), as well as transcripts of 14 tissues (adipose, cerebellum, cerebrum, dorsal muscle, heart, kidney, lung, large intestine, leg muscle, liver, stomach, small intestine, spleen, testis) collected from the assembled Debao pony. Transcripts were obtained through RNA-Seq using HISAT2 (v2.2.1)(*58*) and StringTie (v2.1.4)(*59*), as well as pooled Iso-Seq using IsoSeq3 (v3.4.0, https://github.com/PacificBiosciences/IsoSeq) and minimap2 (v2.20)(*60*). RNA-Seq and Iso-Seq were sequenced on the Illumina NovaSeq 6000 platform and PacBio Sequel platform, respectively. The initial round of annotation utilized GeneMark-

ES (v4.64)(*61*) for gene model training and prediction with evidence support. The second round involved training SNAP (v2006-07-28)(*62*) and AUGUSTUS (v3.4.0)(*63*) with predicted gene models, and the results were integrated to predict gene models. An additional iterative *ab initio* gene prediction step was implemented. Gene models without evidence support were rescued if they were annotated by BUSCO or InterProScan (v5.54-87.0)(*64*). Gene models that were absent compared with the NCBI Annotation Release 103 of EquCab3.0 were merged with our annotation dataset using GFF3toolkit (v2.1.0, https://github.com/NAL-i5K/GFF3toolkit). The final Maker3 gene models were manually curated with spliced alignment data from RNA-Seq and Iso-Seq using Apollo (https://cpt.tamu.edu/galaxy-pub/)(65). The accuracy of the genome annotation was assessed using annotation edit distance (AED). Functional annotation was assigned based on the best hit from DIAMOND (v2.0.14)(*66*) alignment to the TrEMBL database. Motifs and domains of protein-coding genes were determined using InterProScan. GO terms and KEGG pathways were assigned using the best-match classification. Noncoding RNAs were predicted using RNAmmer (v1.2)(*67*), tRNAscan-SE (v2.0.7)(*68*), and Infernal (v1.1.4)(*69*).

### Comparative genomics analysis

The protein-coding genes from 14 genomes, including horse (*Equus caballus*, Thoroughbred & Debao pony), ass (*Equus asinus*), plains zebra (*Equus quagga*), dog (*Canis lupus familiaris*), goat (*Capra hircus*), sheep (*Ovis aries*), cattle (*Bos taurus*), pig (*Sus scrofa*), human (*Homo sapiens*), chimpanzee (*Pan troglodytes*), rhesus monkey (*Macaca mulatta*), house mouse (*Mus musculus*), and platypus (*Ornithorhynchus anatinus*), were employed. The longest protein sequences from alternative transcripts were selected to represent unique genes. Orthologous gene families were inferred using OrthoFinder (v2.5.4)(*70*). Single-copy gene families with more than 100 amino acids were aligned using MUSCLE (v5.1)(*71*), and the resulting protein sequence alignment was used to construct the maximum likelihood (ML) phylogenetic tree with RAxML (v8.2.12)(*72*). Divergence times were estimated using the MCMCTree program of PAML (v4.9)(*73*) with TimeTree database (http://www.timetree.org/). Gene family expansion and contraction were detected using CAFE (v5.0)(*74*). GO enrichment analysis was performed using the R package clusterProfiler (v4.6.0)(*75*).

### Resequencing and variant calling

Blood samples were collected from 10 Debao ponies, 10 Baise horses, and 6 Warmbloods. Genomic DNA was extracted using the DNeasy Blood & Tissue Kit (Qiagen, Beijing, China), and DNA integrity was assessed using agarose gels. PE sequencing libraries were prepared and sequenced on an Illumina NovaSeq 6000 platform, generating 150-bp reads with a target depth of 10× coverage. Publicly available whole-genome sequencing (WGS) data for 376 domestic horses, eight Przewalski’s horses, five domestic donkeys, and one Somali wild ass (*E.a.som*) were downloaded from NCBI. Detailed information for all 416 individuals is provided in table S4.

Raw WGS data underwent quality control using fastp. Clean reads were mapped to the assembled Debao pony genome using BWA-MEM (v0.7.17-r1188)(*76*) with default parameters (table S6). The mapped reads were sorted and converted to BAM format using SAMtools (v1.12)(*77*). Variations were detected using the GATK (v4.2.0.0) germline short variant discovery pipeline (https://gatk.broadinstitute.org/). In brief, duplicate reads were identified using the MarkDuplicates module. Variations were called via local re-assembly of haplotypes using the HaplotypeCaller module. Raw genomic variant call format (GVCF) files were merged using the GenomicsDBImport module, and joint genotyping was performed using the GenotypeGVCFs module. The resulting variant calls (SNPs and Indels) were subjected to hard-filtering based on specific parameter distributions: QD < 2.0 || MQ < 50.0 || FS > 60.0 || SOR > 3.0 || MQRankSum < −5.0 || ReadPosRankSum < −5.0 || QUAL < 30.0 (fig. S10). SNP dataset was annotated using SnpEff (v5.1d)(*78*).

### Population genetics analysis

Population genetic analysis was conducted using a dataset of horse samples (table S4). To ensure the reliability of breed records and the purity of lineages, potential crossbred samples were excluded based on population structure inconsistencies. Filtering of SNPs was performed using PLINK (v1.90b6.21)(*79*) with the following criteria: missing genotype rate per sample > 30%, missing genotype rate per variants > 20%, minor allele frequency (MAF) < 0.01, Hardy-Weinberg equilibrium < 0.000001, and SNPs pruned in LD block (window size: 50; step size: 5; r^2^ threshold: 0.5). The resulting dataset consisted of 4,881,874 autosomal SNPs from 385 samples representing 49 horse breeds, which were used for subsequent analyses. PCA was performed using GCTA (v1.93.2beta)(*80*) with default settings. For phylogenetic analysis, the p-distance matrix was calculated using VCF2Dis (https://github.com/BGI-shenzhen/VCF2Dis). Based on the p-distance matrix, a neighbor-joining (NJ) phylogenetic tree was constructed using the fneighbor command of EMBOSS (v6.6.0.0)(*81*). The tree was rooted on Przewalski’s horses and visualized using the R package ggtree (v3.6.2)(*82*). Ancestry and population structure were analyzed using ADMIXTURE (v1.3.0)(*83*) with *K* values ranging from 2 to 20. In the SNPs pruning step, the parameters used were window size: 50, step size: 10, and r^2^ threshold: 0.1. The optimal *K* value was determined based on the cross-validation error.

### Y chromosome haplotype

Y chromosome analysis was conducted on male individuals (table S4). The sex of each individual was determined based on the read coverage on the Y chromosome, resulting in the identification of 177 male horses with read coverage exceeding 93.9% on the Y chromosome (fig. S13 and table S11). SNPs specific to the Y chromosome were extracted, and heterozygous sites were excluded to minimize false positive rates in the male-specific region of the Y chromosome (MSY). The remaining SNPs were filtered using specific criteria: missing genotype rate per sample > 20% and missing genotype rate per variant > 10%. A phylogenetic tree rooted on Przewalski’s horses was constructed based on the p-distance matrix calculated using VCF2Dis and the fneighbor command of EMBOSS. For haplotype analysis, variant sites with missing genotypes were removed. Haplotypes were identified, and a haplotype network was computed using the R package pegas (v1.2)(*84*).

### Genome evolution analysis

Genome evolution analysis included a dataset of horse and donkey samples (table S4)(*85*). SNPs were filtered using PLINK with the following criteria: missing genotype rate per sample > 30%, missing genotype rate per variants > 20%, minor allele frequency (MAF) < 0.005, Hardy-Weinberg equilibrium < 0.000001, and SNPs pruned in LD block (window size: 50; step size: 5; r^2^ threshold: 0.5).

Historical population relationships were inferred by constructing the ML phylogenetic tree and calculating outgroup *f3*-statistics. The ML tree was built using TreeMix (v1.13)(*86*) with variants present in all individuals. SNPs were further pruned with specific parameters: window size: 50, step size: 10, and r^2^ threshold: 0.2. This resulted in a dataset of 15,497,130 autosomal SNPs, which were transformed into TreeMix input format using the plink2treemix.py script. The ML tree was rooted on *E.a.som* with 1000 bootstrap replicates. Shared genetic drift between pairs of populations was quantified by calculating the *f3*-statistics in the form *f3*(X, Y, outgroup), where X and Y represent all possible pairwise combinations of domestic horse breeds included in this study, and outgroup represents *E.a.som*. The *f3*-statistics were computed with 59,971,670 autosomal SNPs using the R package ADMIXTOOLS 2 (v2.0.0)(*87*).

LD decay patterns were measured in two groups (tables S4 and S5): 106 top-height horses (Shire horse, Percheron, Oldenburger, Holsteiner, Württemberger, Hanoverian horse, Dutch Warmblood, Westphalian horse, Bavarian Warmblood, Warmblood, Trakehner, Thoroughbred, and Standardbred) and 93 bottom-height horses (Miniature horse, Debao pony, Shetland pony, Baise horse, Jeju horse, Tibetan pony, Erlunchun horse, Dülmener pony, and Chakouyi horse). The LD decay analysis was performed using PopLDdecay (v3.42)(*88*).

A two-dimensional PHATE embedding was generated using the top 98 principal components from the PCA as input in the R package phateR (v1.0.7)(*89*), which exhibited a clear hierarchical structure and transition state trajectories. The parameters used were: ndim = 2, knn = 5, decay = 40, gamma = 1, t = 26, and seed = 1234.

Gene flows of pony populations were inferred with TreeMix using representative populations (Przewalski’s horse, Icelandic horse, Baise horse, Debao pony, Yakutian horse, Tibetan pony, Jeju horse, Mongolian horse, Akhal-Teke, Arabian horse, and Thoroughbred). The SNPs were filtered as described in the above TreeMix procedure. The topology was rooted on *E.a.som* with 1000 bootstrap replicates. The TreeMix model was run from 0 to 30 migration events with 10 replicates. The optimal migration edges were inferred using the R package optM (v0.1.6) with 1000 bootstraps performed using the R package BITE (v1.2.0008)(*90*).

The *f4*-statistics were calculated to assess the genetic relationship between populations with autosomal SNPs used in the *f3*-statistics analysis. The *f4*-statistics were modelled as *f4*(P1, P2, P3, OG) using the R package ADMIXTOOLS 2. Where P1, P2, and P3 represent all possible permutations of domestic horse breeds, and OG represents *E.a.som* as the outgroup. A relationship network was constructed by integrating every possible combination of *f4*-statistics, and distances between edges were calculated using the following formulas:

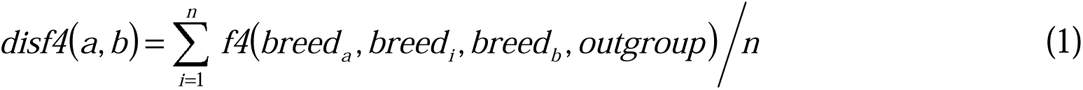

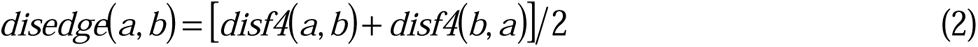

Where *disf4*(*a*, *b*) represents the directional distance from *breed_a_* to *breed_b_*; *f4*(*a*, *i*, *b*, *outgroup*) represents the specific *f4*-statistics formulation; *n* represents the number of horse breeds included in the analysis; *disedge*(*a*, *b*) represents the non-directional distance between *breed_a_* and *breed_b_*.

Demographic history of representative horse populations (Debao pony, Erlunchun horse, Yakutian horse, Connemara pony, and Shetland pony) in the relationship network was estimated using SMC++ (v1.15.5)(*91*). A mutation rate of 7.242C×C10^-9^ per base pair per generation and a generation time of eight years were assumed to convert coalescence generations into demographic events of *N_e_*.

### Identification of height-related genes

Identification of height-related genes involved a comprehensive analysis of horse and pony samples (tables S4 and S5). The dataset included 26 horse individuals representing top height breeds (Shire horse, Percheron, Oldenburger, Holsteiner, Württemberger, Hanoverian horse, Dutch Warmblood, Westphalian horse, Bavarian Warmblood, Warmblood, and Trakehner) and 31 pony individuals representing bottom height breeds (Shetland pony, Debao pony, and Miniature horse).

The variant dataset was imputed and phased using Beagle (v5.4)(*92*). To detect selective signals, scans for selection signatures were performed using fixation index (*F*_ST_), nucleotide diversity (π) ratio (θπ_pnoy_/θπ_horse_), and cross-population extended haplotype homozygosity (XP-EHH). Genetic statistics of *F*_ST_ and θπ were calculated across the whole genome using 10 kb nonoverlapping sliding windows with VCFtools (v0.1.16)(*93*). The XP-EHH analysis was performed with bi-allelic sites using selscan (v1.2.0)(*94*), and averaged XP-EHH scores were calculated with 10 kb nonoverlapping sliding windows. Genes located in genomic regions with top 5% values in all three tests were subjected to KEGG pathway enrichment analysis using the R package clusterProfiler. Gene functions were annotated using eggNOG-mapper (v2.1.8)(*95*), and the organism package for DeBao1.0 assembly was built using the R package AnnotationForge (v1.40.0). Genomic regions with top 1% values in at least two selection signatures were considered selective sweeps. Only genes in the selective sweep regions with potential impacts on body size were identified as candidate height-related genes.

Hi-C data analysis was conducted using the HiCExplorer (v3.7.2) pipeline(*96*). To ensure a robust result, Hi-C data from both the Debao assembly in this study and the EquCab3.0 assembly were merged initially. Hi-C reads preprocessed by fastp were mapped to the DeBao1.0 genome using BWA-MEM with specific parameters (-A1, −B4, −E50, and −L0). The aligned reads in BAM format were filtered, and a contact matrix with a bin size of 10 kb was created using the hicBuildMatrix function. The Hi-C matrix was then normalized and corrected with a threshold of 1.5 using the hicNormalize and hicCorrectMatrix functions, respectively. Topologically associating domains (TADs) were identified using the hicFindTADs function.

Regulation elements were annotated utilizing ChIP-Seq data obtained from the FAANG database (https://data.faang.org). Specifically, histone marks including H3K4me1, H3K4me3, H3K27ac, and H3K27me3 were retrieved from the diaphysis of metacarpal bone and then processed using Trim Galore (v0.6.7, https://www.bioinformatics.babraham.ac.uk/projects/trim_galore/). Clean reads were subsequently mapped to the DeBao1.0 genome using Bowtie 2 (v2.4.5)(*97*) with default parameters. To ensure data quality, non-aligned and poorly mapped reads (MAPQ < 10) were filtered out using SAMtools. The aligned reads were sorted, and duplicate alignments were marked using SAMtools. ChIP-Seq peak calling was performed employing MACS2 (v2.2.7.1)(*98*). Bigwig files were generated for track visualization using deepTools (v3.5.1)(*99*).

Runs of homozygosity (ROH) were analyzed using the --homozyg function in PLINK with specific parameters (table S4): ROH ratio (kb/variant) ≤ 50, between-variant distance within an ROH ≤ 100, ROH length ≥ 500 kb, ROH variant count ≥ 50, hets in a scanning window hit ≤ 1, scanning window size (SNP) = 50, and a scanning window hit rate for a variant to be included in an ROH ≥ 0.05 (fig. S18).

PhyloP scores were calculated based on conservation scoring by phyloP (phylogenetic p-values)(*100*) for multiple alignments of placental mammal to the human genome, which were available at (http://hgdownload.cse.ucsc.edu/goldenPath/hg19/phyloP46way/placentalMammals/). Briefly, the alignment was achieved by aligning the DeBao1.0 genome to the EquCab2.0 genome using minimap2. Subsequently, a chain file was created from minimap2 using transanno (v0.3.0, https://github.com/informationsea/transanno). Genome coordinates were then converted with the chain file created above and obtained from UCSC (https://hgdownload.soe.ucsc.edu/goldenPath/equCab2/liftOver/equCab2ToHg19.over.chain.g <u>z</u>) using liftOver (v447)(*101*). Lastly, blocks with a distance of less than 10 bp were merged using BEDTools (v2.30.0)(*102*).

Candidate regions were visualized using the R package plotgardener (v1.4.2)(*103*). For most of the visualizations, the R package ggplot2 (v3.4.2) was utilized unless otherwise specified.

## Acknowledgments

We thank Hongru Wang for comments and critical reading of the manuscript.

## Funding

Guangxi special project for innovation-driven development (AA17204024)

Agricultural Science and Technology Innovation Program of the Chinese Academy of Agricultural Sciences (CAAS-ASTIP)

## Author contributions

Conceptualization: G.Y., Y.L., Z.T., X.L.

Investigation: Z.W., M.Z., B.W., X.Q., M.C.

Resources: S.T., J.Y., S.C., P.Y.

Data curation: X.L., Q.B., S.H.

Methodology: X.L., Z.W., X.Y.

Formal Analysis: X.L., L.L., P.Z. Visualization: X.L.

Supervision: G.Y., Y.L., Z.T. Writing—original draft: X.L.

Writing—review & editing: G.Y., L.F., H.X., J.J., X.L. Funding acquisition: G.Y., Y.L.

## Data and materials availability

The genome assembly and raw sequencing data generated in this study, including PacBio CLR data, Hi-C data, PE sequencing data, Iso-Seq data, RNA-Seq data, and NGS data have been deposited in the NCBI database under BioProject accession PRJNA1005486. The detail runs of downloaded NGS data from NCBI SRA database are shown in table S4. All data needed to evaluate the conclusions in the paper are present in the paper and/or the Supplementary Materials.

