## Supplementary information for "Genomic insights into post-domestication expansion and selection of body size in ponies"

Xingzheng Li *et al.*

**This PDF file includes:**

Figs. S1 to S22

Tables S1 to S18

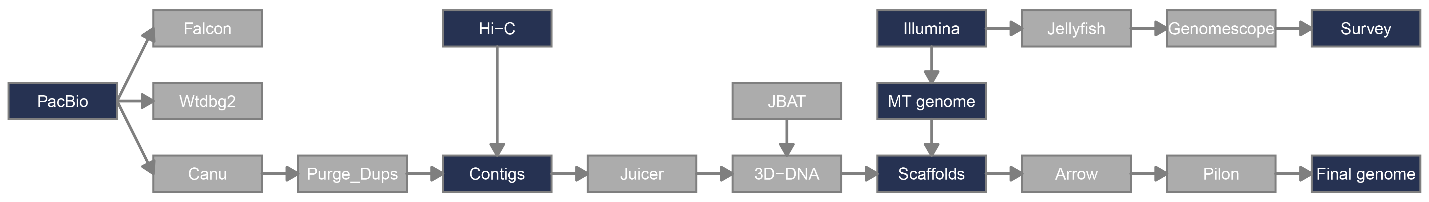

Fig. S1. Flow chart of the genome assembly pipeline used to generate DeBao1.0 assemblies in this study. Dark rectangles represent raw input data or key output.

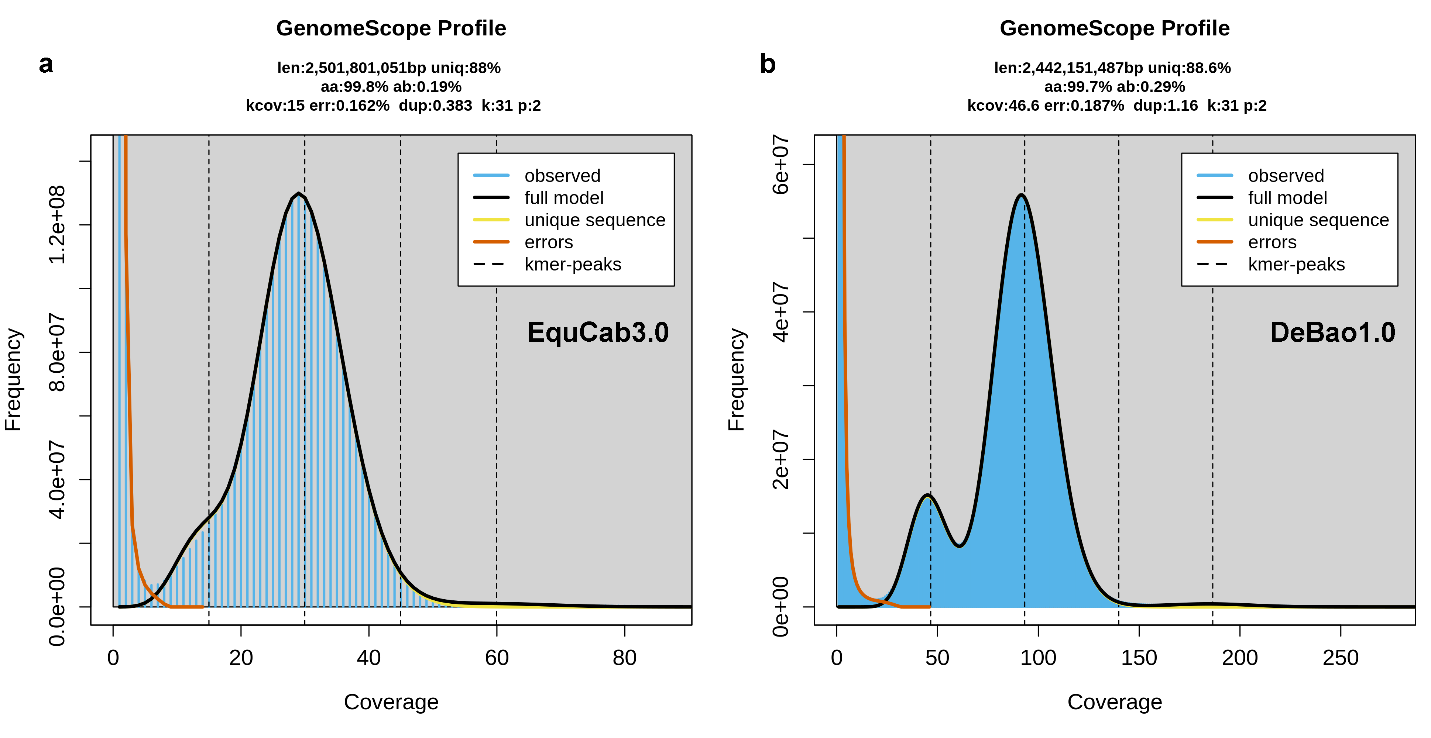

Fig. S2. GenomeScope profiles generated from Illumina short read data for *Equus caballus*. The figure presents the k-mer spectra and their corresponding fitted models for (a) the EquCab3.0 assembly and (b) the DeBao1.0 assembly.

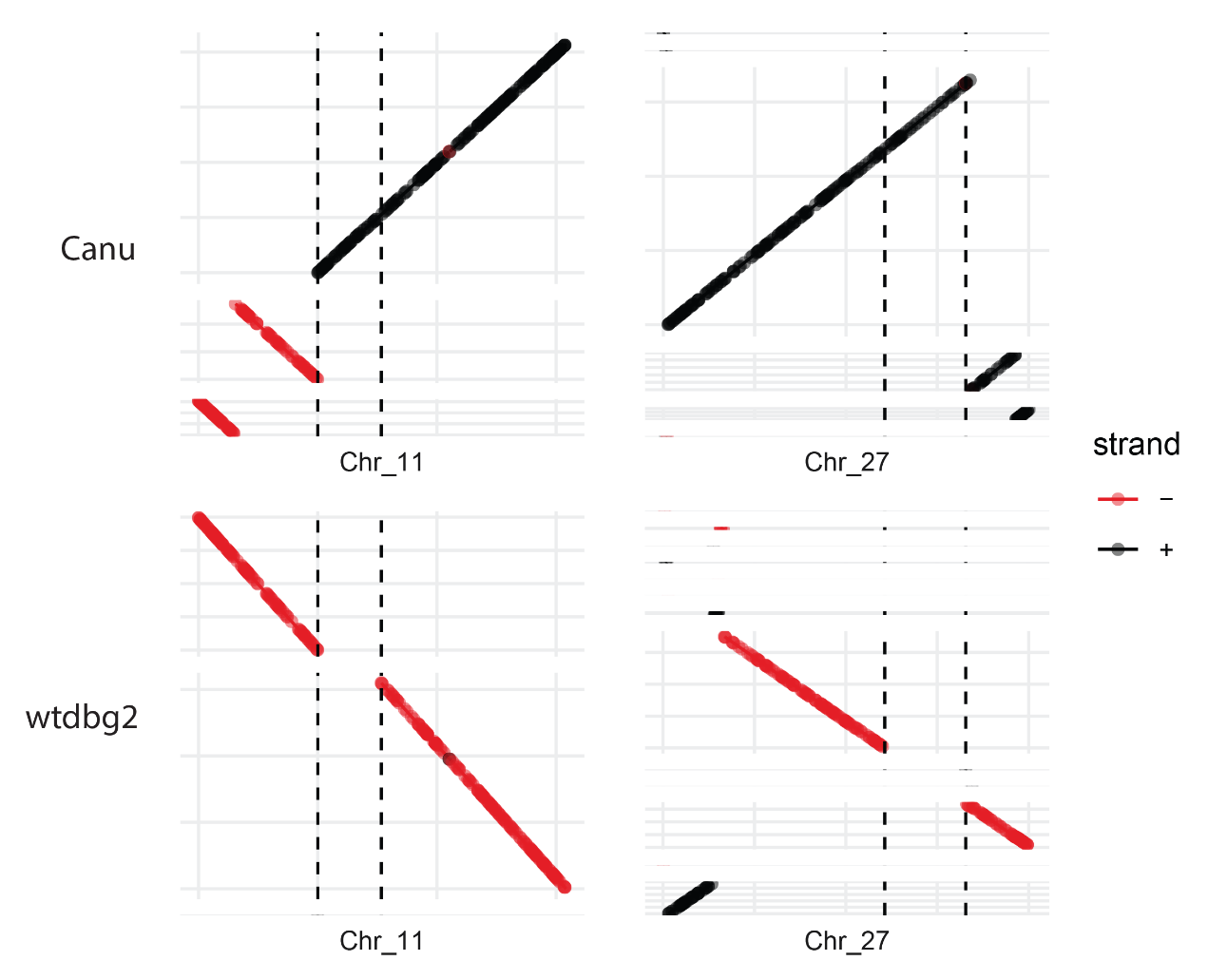

Fig. S3. Illustration of primary assembly completeness achieved by Canu and wtdbg2. The primary assemblies were aligned to the EquCab3.0 genome using MUMmer. The x-axis represents EquCab3.0, and the y-axis represents primary contigs. Missing sequences in the wtdbg2 assembly are indicated between black dashed lines.

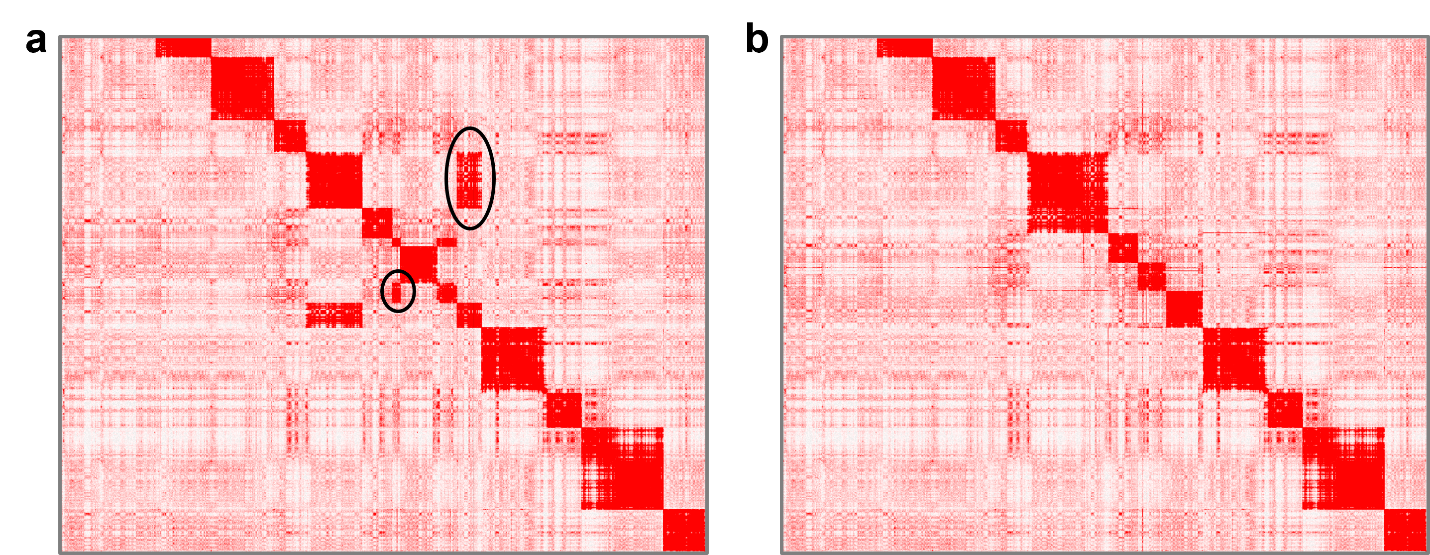

Fig. S4. Demonstration of misassembly correction using Hi-C data. (a) Hi-C map illustrating the pony assembly obtained by scaffolding without any editing, with misjoins indicated by black ellipses. (b) Hi-C map after the misjoins have been corrected.

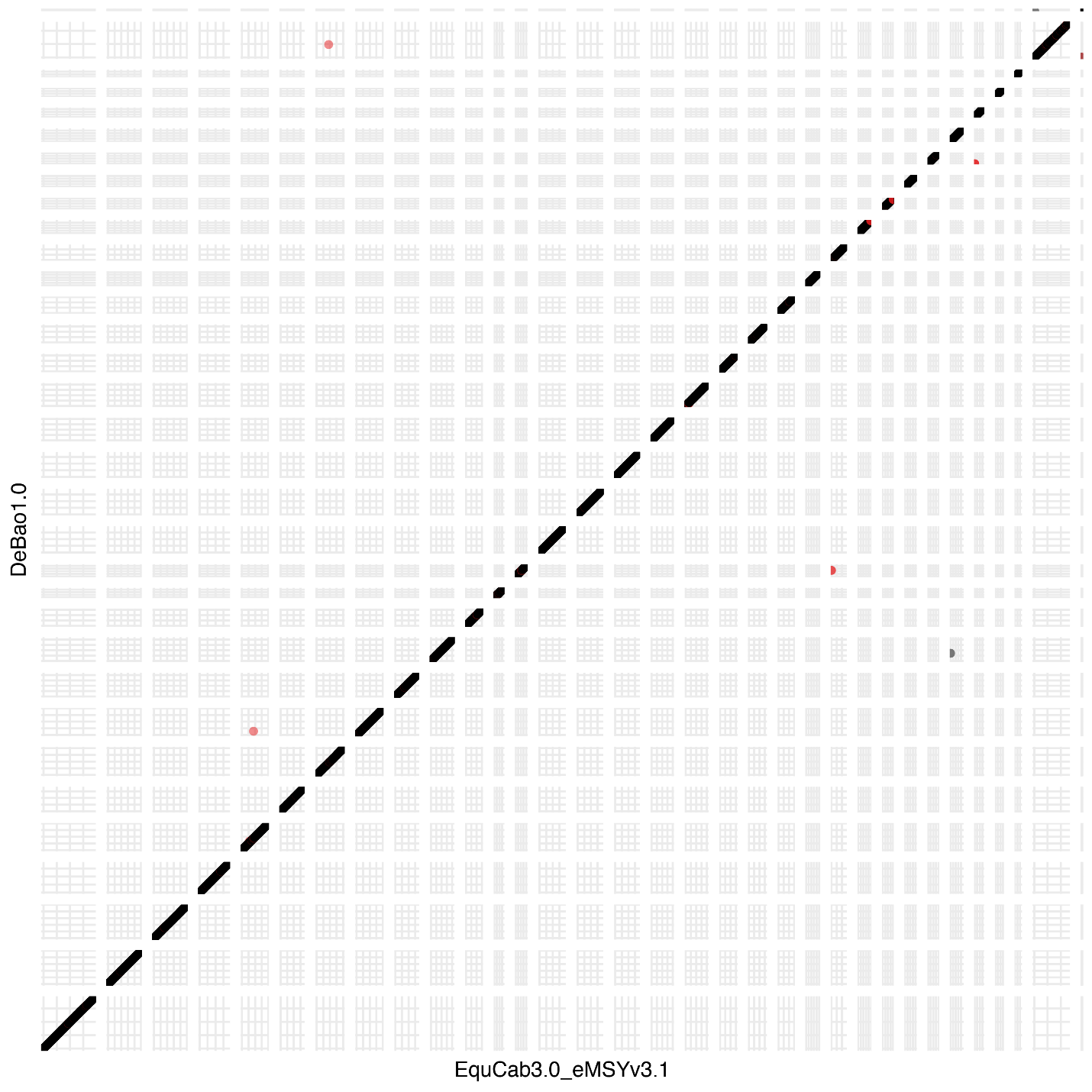

Fig. S5. Synteny between DeBao1.0 and EquCab3.0 chromosomes. The Y chromosome assembly (eMSYv3.1) is appended to EquCab3.0. The black color represents alignment to the forward strand, and the red color represents alignment to the reverse strand.

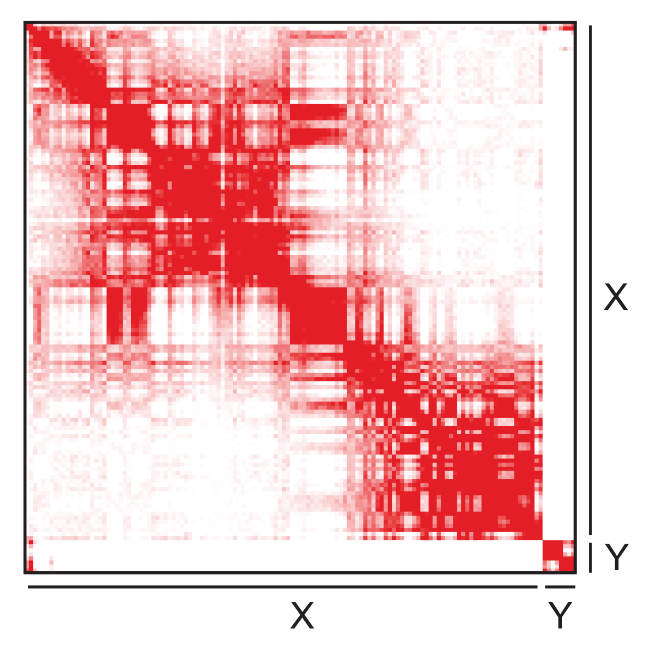

Fig. S6. Hi-C chromatin interactions of the assembled sex chromosome.

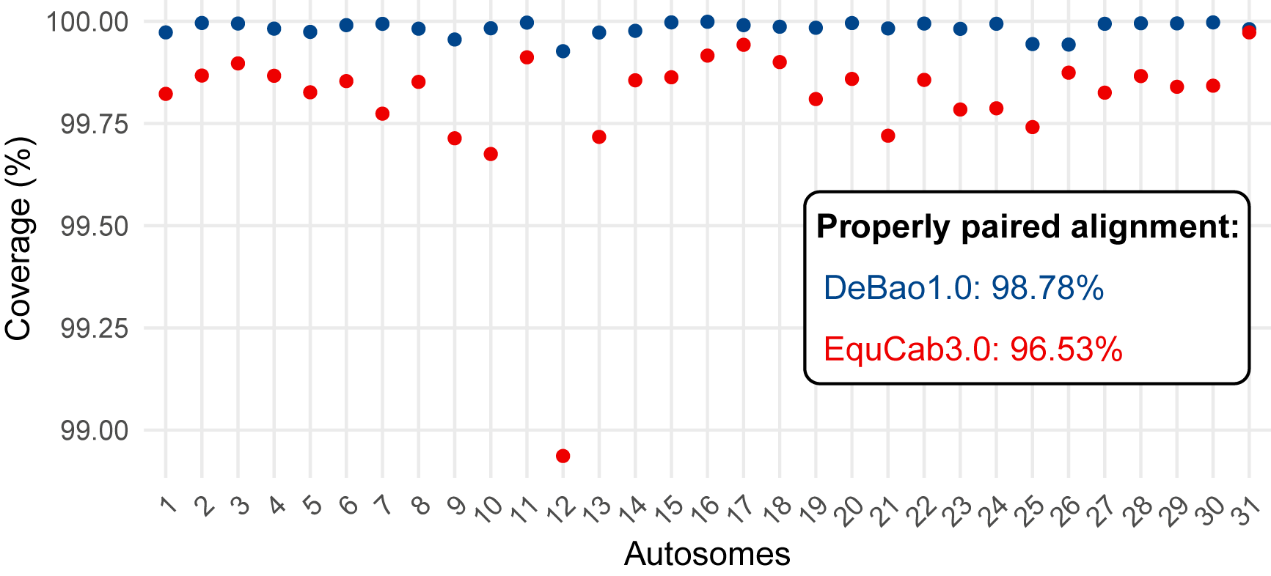

Fig. S7. Alignment statistics of read coverage per autosome and the overall alignment rate for the DeBao1.0 and EquCab3.0 genome assemblies. Illumina paired-end (PE) reads involved in generating each corresponding assembly were utilized for calculating these statistics.

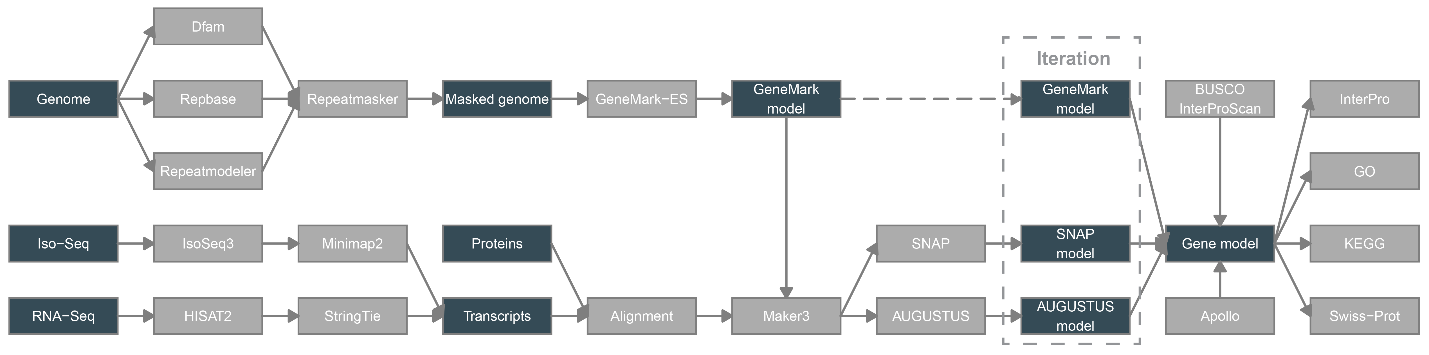

Fig. S8. Flow chart of the genome annotation pipeline used to annotate DeBao1.0 assembly in this study. Dark rectangles represent raw input data or key output.

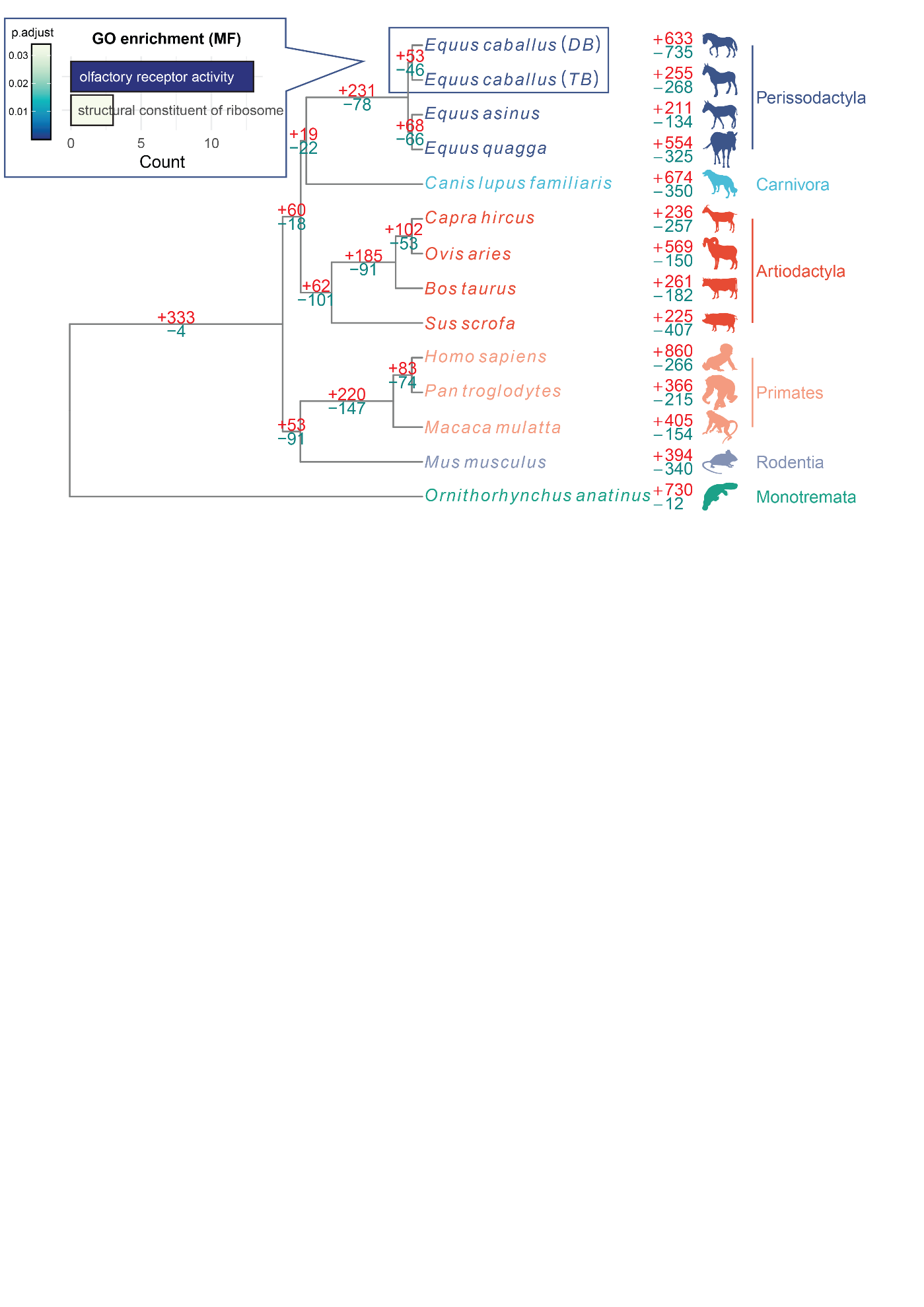

Fig. S9. Gene family evolution of *Equus caballus* in a total of 13 species. The number of gene gains and losses is indicated at the corresponding nodes and branches. The Gene Ontology (GO) molecular function (MF) enrichment analysis for the contracted gene family of *Equus caballus* is presented at the top left. *Equus caballus* (*DB*) represents the DeBao1.0 assembly of the Debao pony, and *Equus caballus* (*TB*) represents the EquCab3.0 of the Thoroughbred.

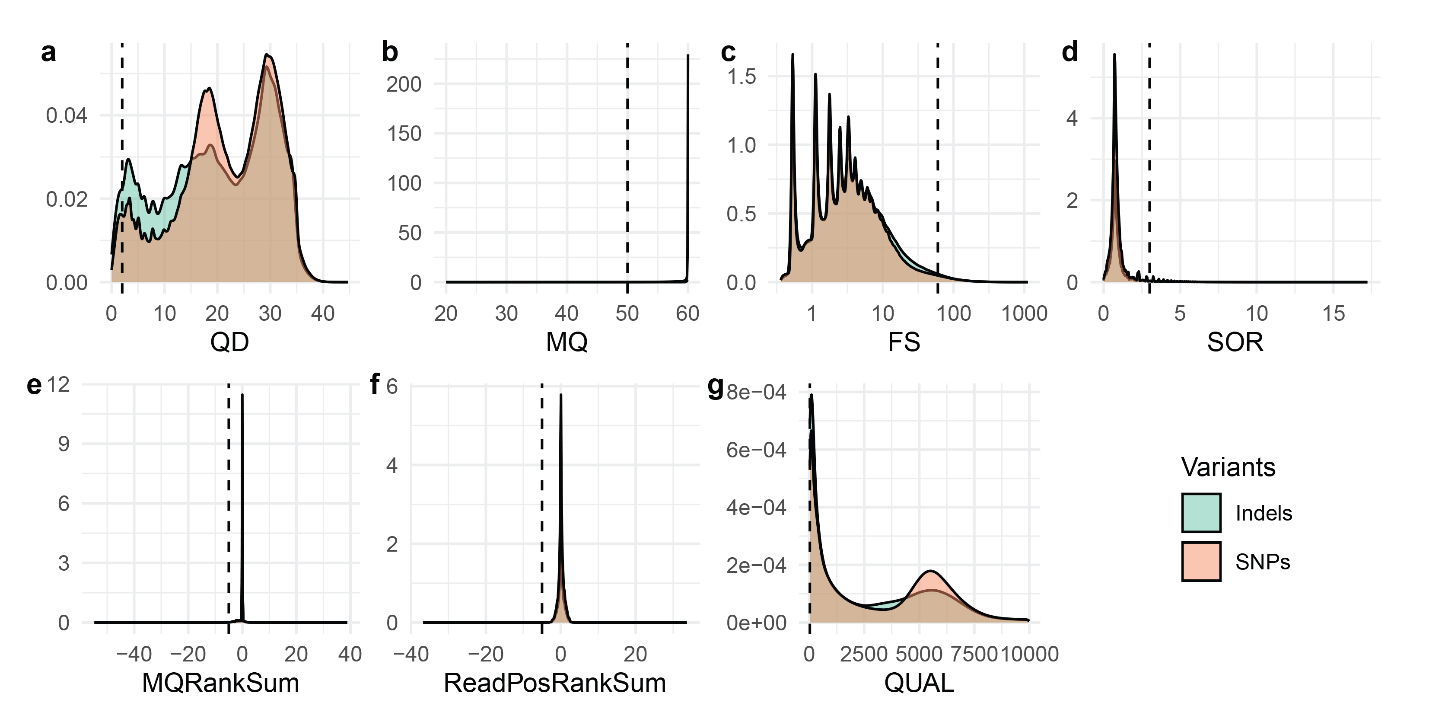

Fig. S10. Distribution of hard-filtering parameters in GATK variant calling (SNPs & Indels). These parameters include (a) QualByDepth (QD), (b) RMSMappingQuality (MQ), (c) FisherStrand (FS), (d) StrandOddsRatio (SOR), (e) MappingQualityRankSumTest (MQRankSum), (f) ReadPosRankSumTest (ReadPosRankSum), and (g) QUAL. The parameters used are indicated with black dashed lines.

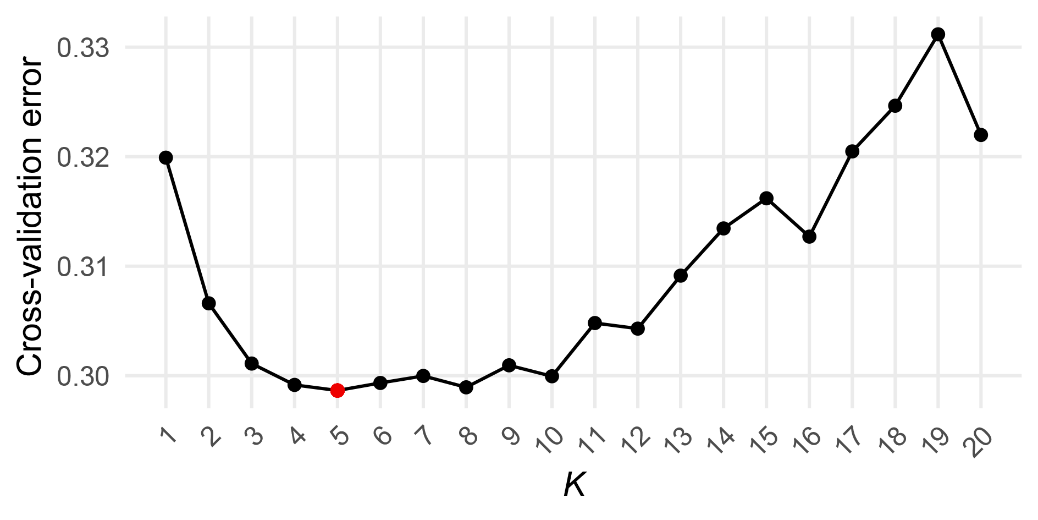

Fig. S11. Cross-validation errors of ADMIXTURE with *K* values ranging from 1 to 20. The optimal *K* value is indicated in red.

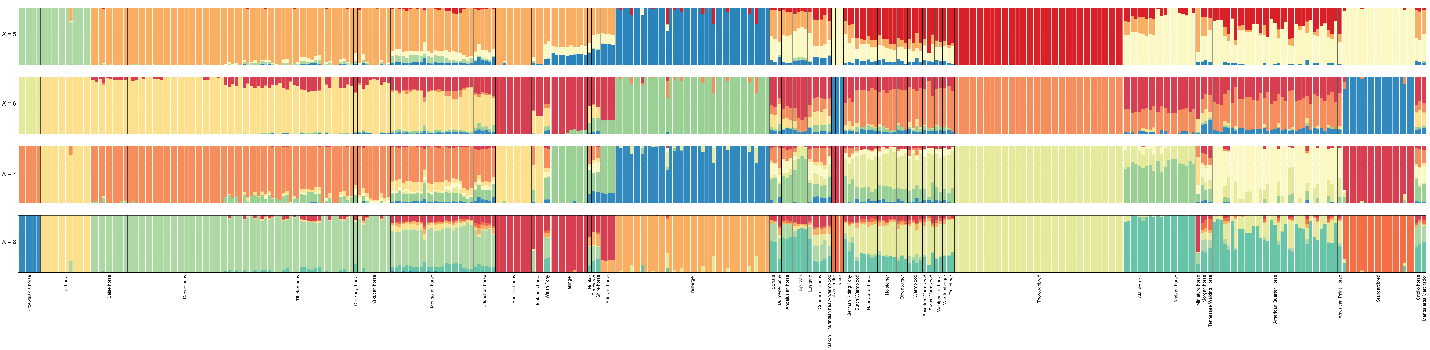

Fig. S12. Admixture analysis of the horse population with *K* values ranging from 5 to 8.

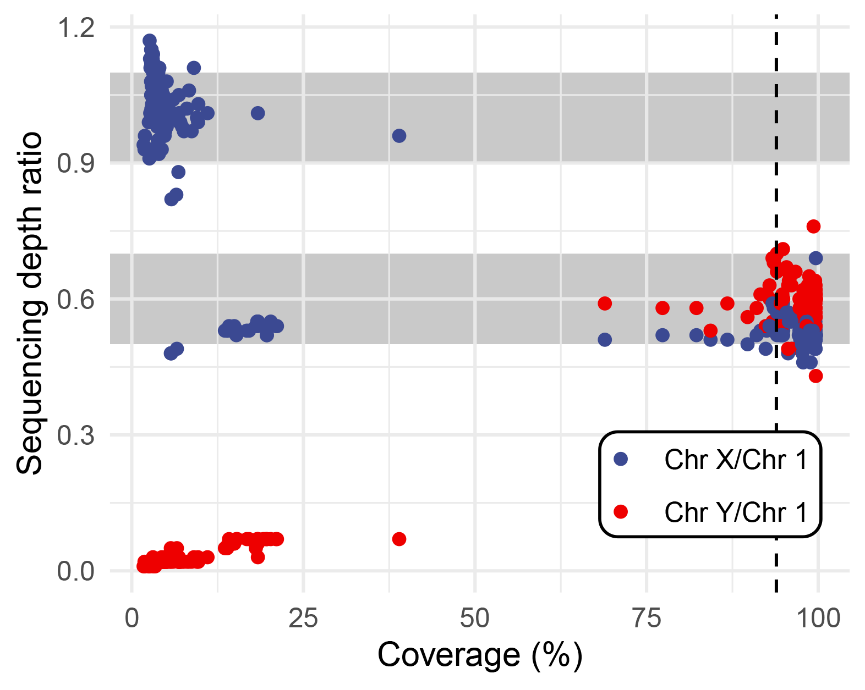

Fig. S13. Read coverage vs. sequencing depth on the Y chromosome. The sequencing depth of chromosome 1 is calculated to represent the overall genome sequencing depth. The grey rectangles, from top to bottom, represent the approximate sequencing depth ratio (Chromosome X/Chromosome 1) ranges corresponding to female and male individuals, respectively. The threshold of read coverage used to determine male individuals is indicated with a black dashed line. Each point represents a horse individual.

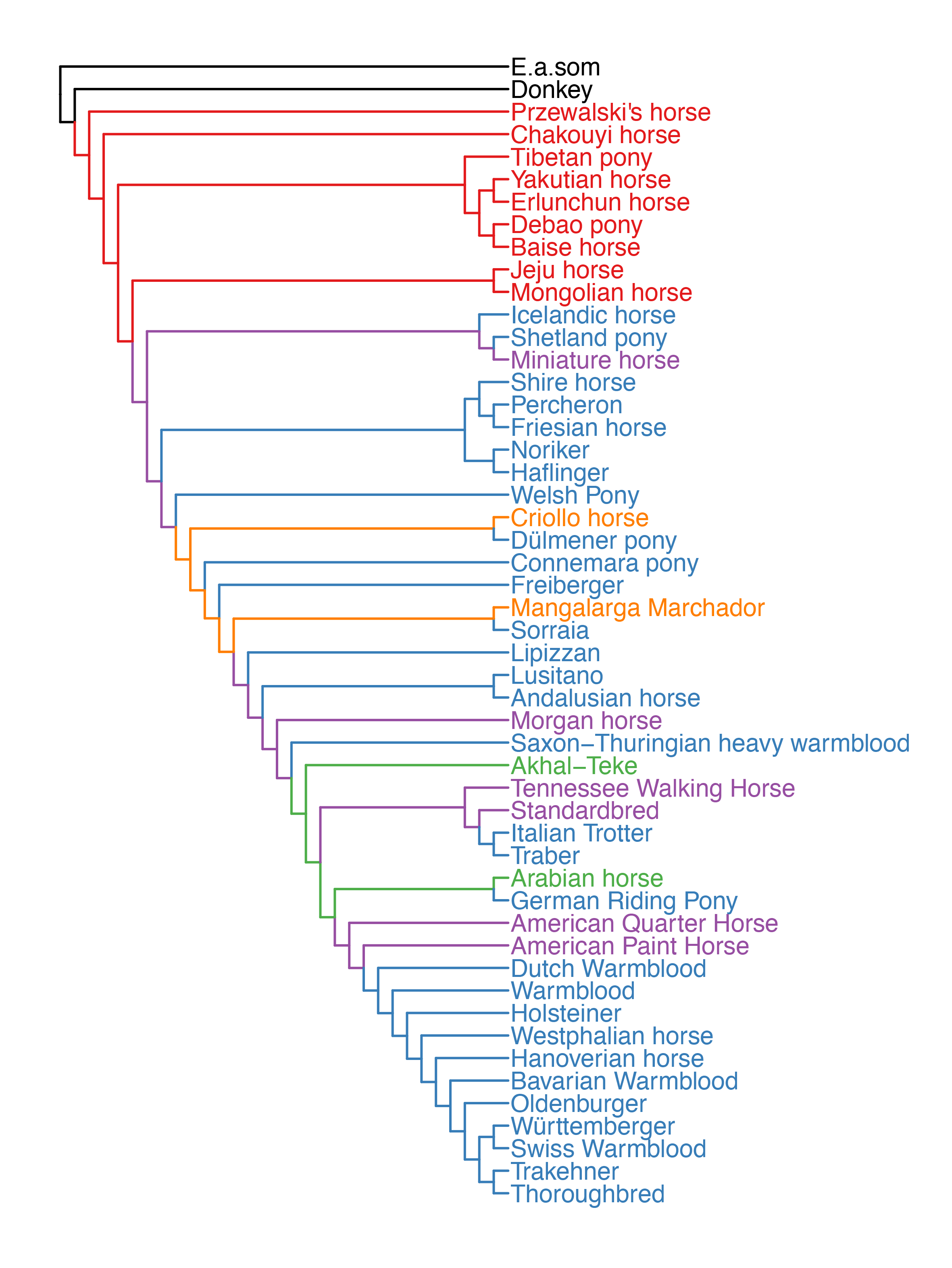

Fig. S14. Phylogenetic relationship reconstruction using TreeMix. This figure corresponds to Fig. 4A.

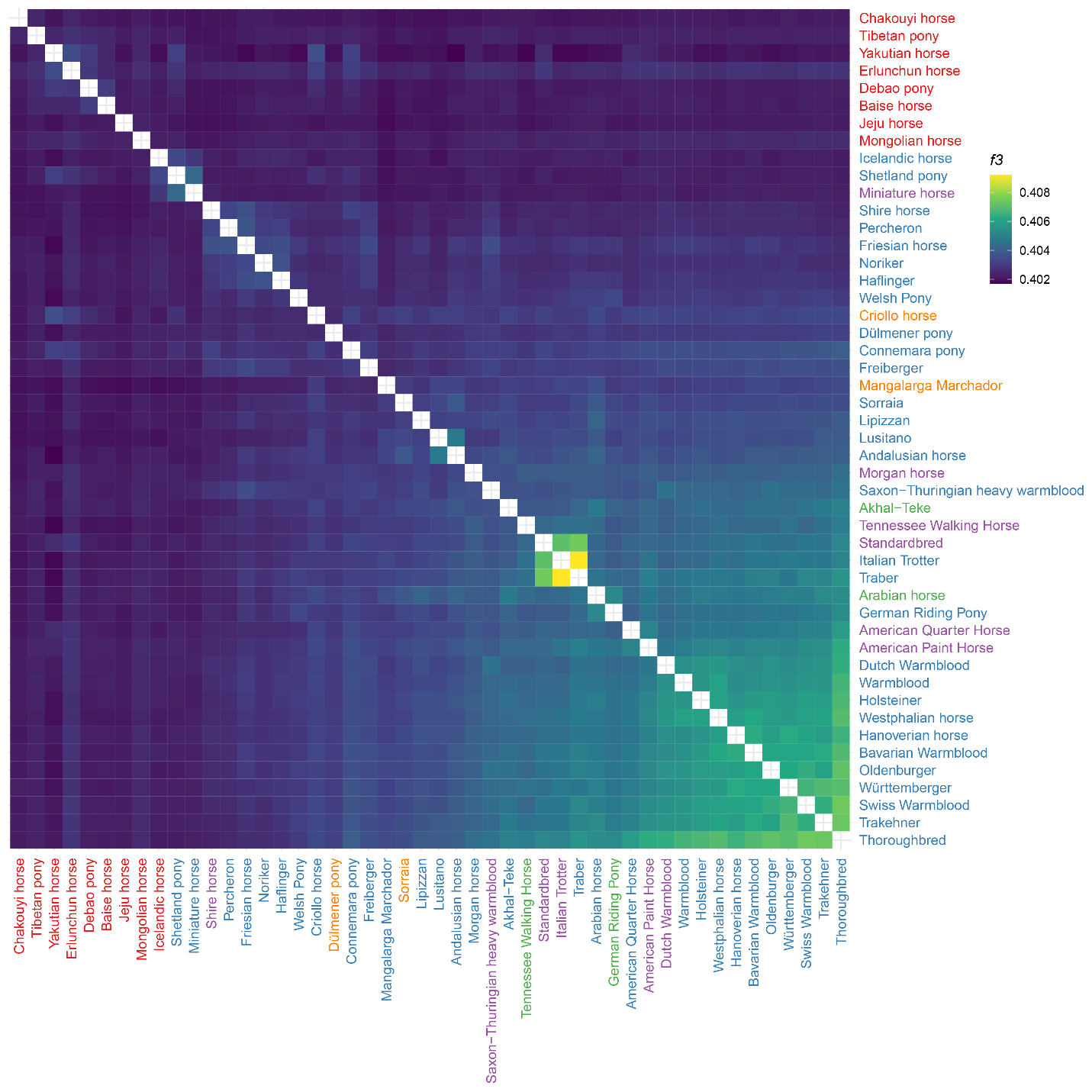

Fig. S15. Heatmap displaying outgroup *f3*-statistics in the form of (*Equus caballus*, *Equus caballus*, *E.a.som*). This figure corresponds to Fig. 4A.

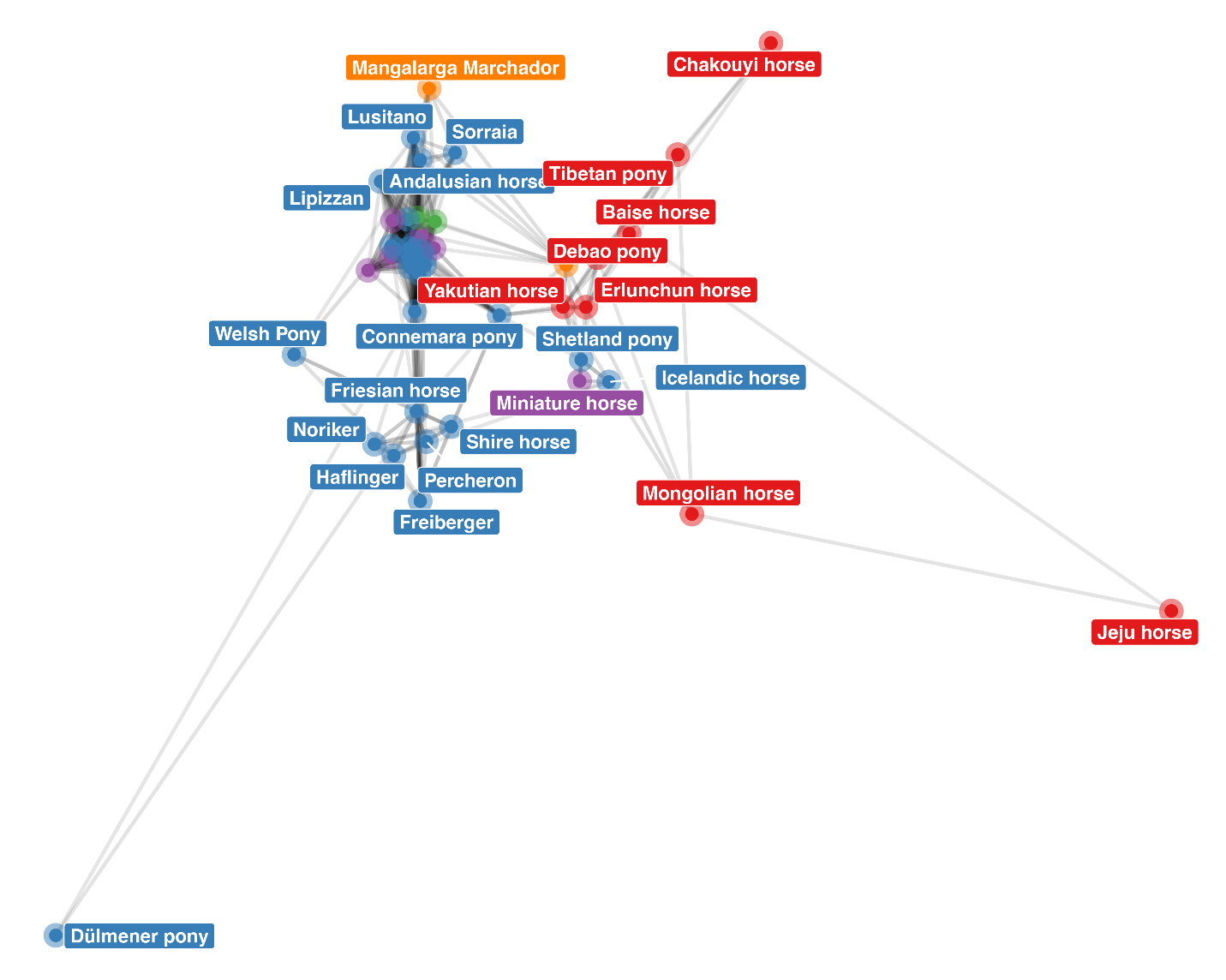

Fig. S16. Relationship network based on every permutation of *f4*-statistics in the form of (*Equus caballus, Equus caballus, Equus caballus, E.a.som*). Line colors ranging from light to dark indicate genetic relationships from distant to close. The pony-related network shown in Fig. 4e is derived from this network.

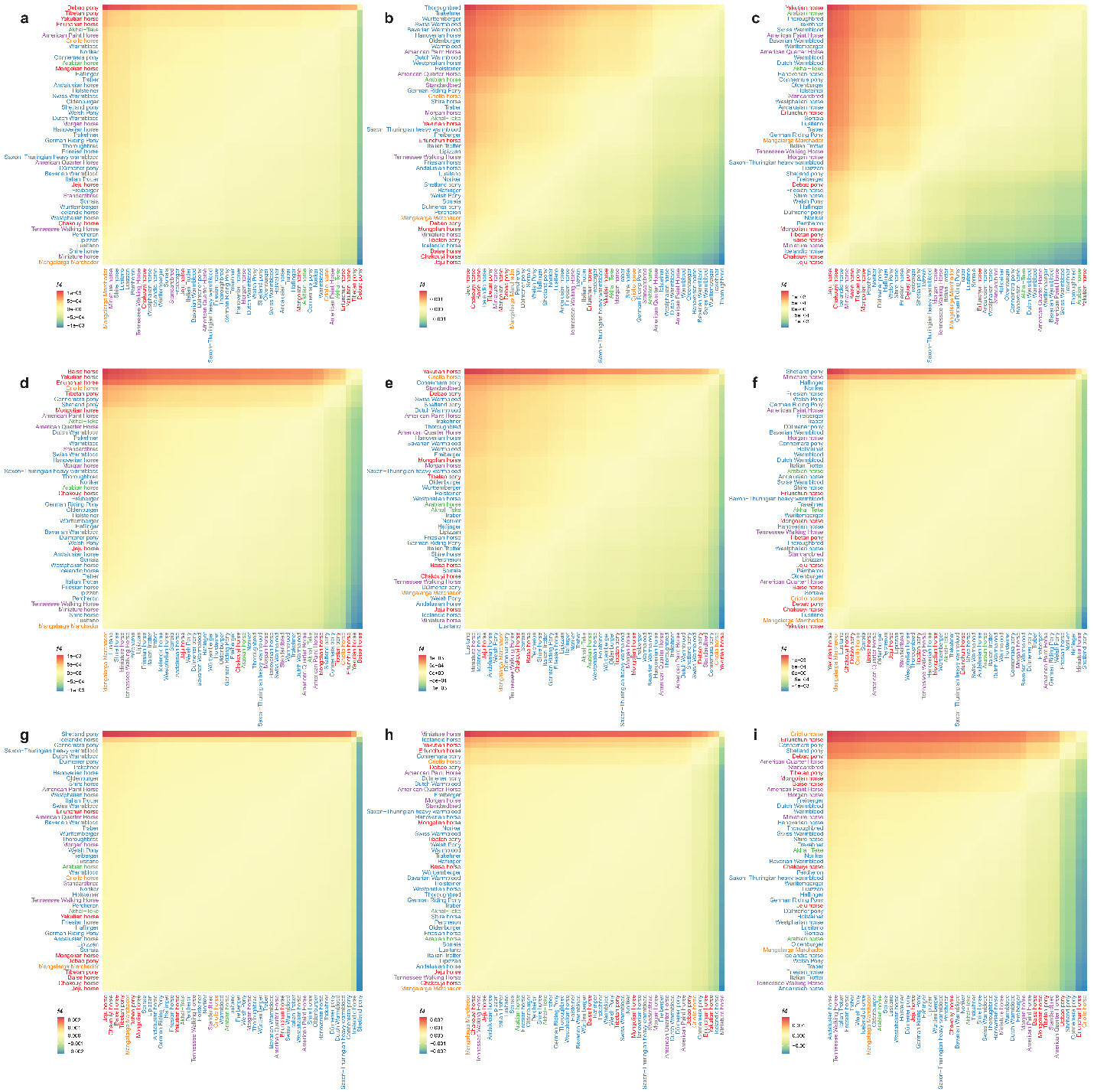

Fig. S17. Heatmap displaying *f4*(P1*_Equus caballus_,* P2*_Equus caballus_,* P3*_Equus caballus_,* OG*_E.a.som_*) for key nodes in the pony-relationship network. This figure corresponds to the nodes in Fig. 4E, with (a) Baise horse, (b) Connemara pony, (c) Criollo horse, (d) Debao pony, (e) Erlunchun horse, (f) Icelandic horse, (g) Miniature horse, (h) Shetland pony, and (i) Yakutian horse as P3. Each row represents P1, and each column represents P2.

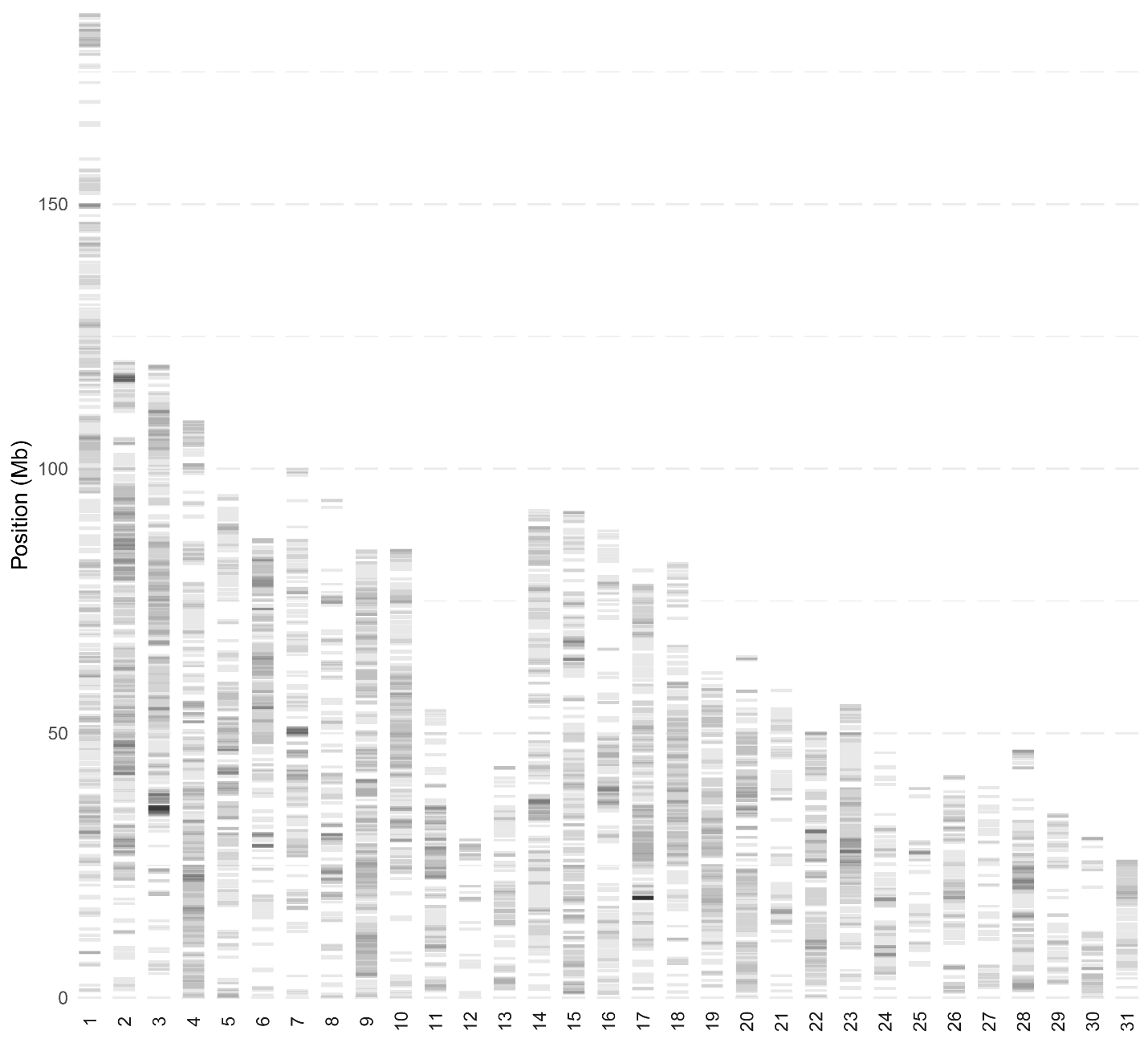

Fig. S18. Genome-wide distribution of runs of homozygosity (ROH) across autosomes in pony-sized populations. The degree of darkness is proportional to the number of individuals with ROH.

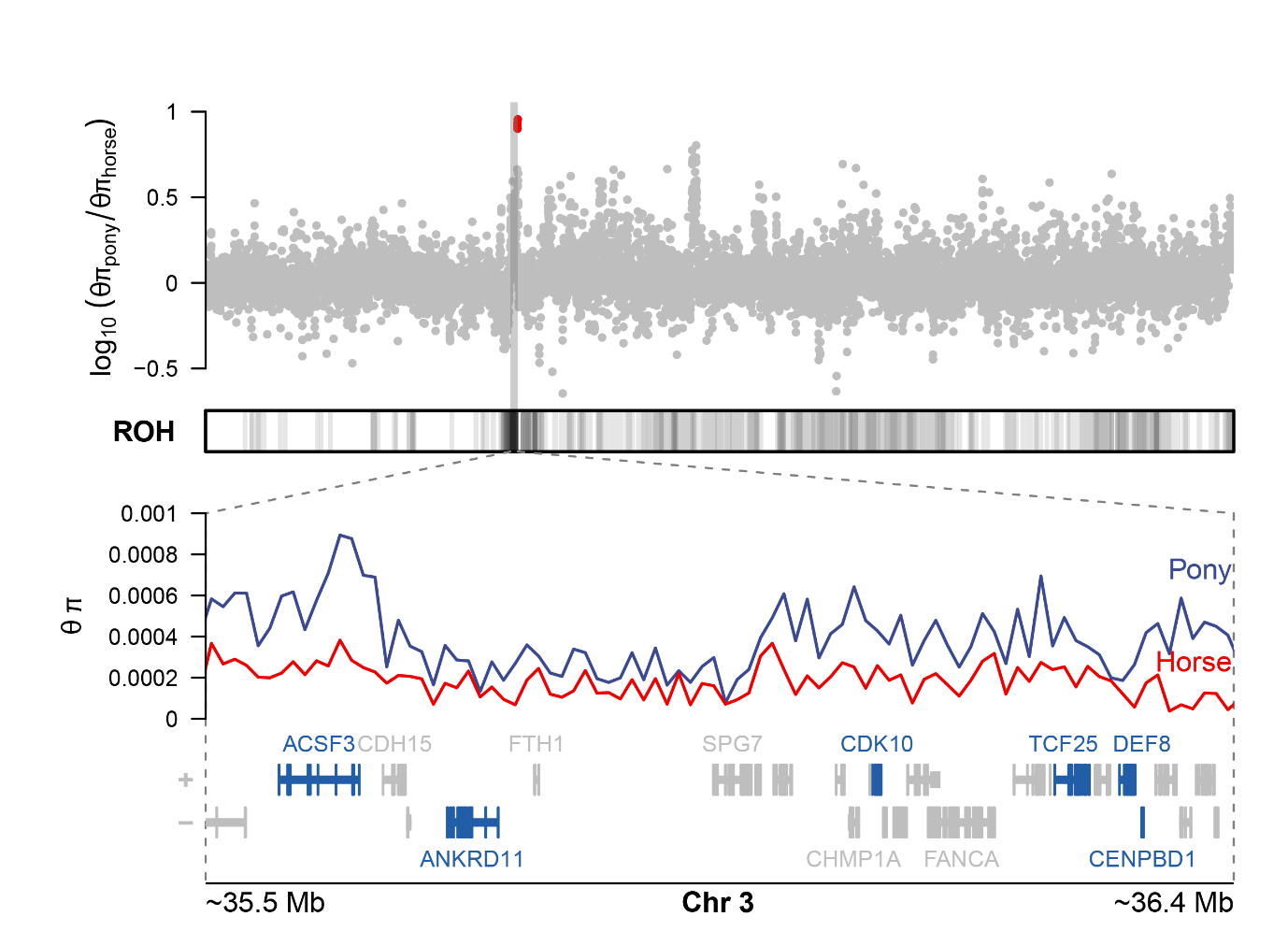

Fig. S19. Candidate selective region on chromosome 3. From top to bottom are π ratio (θπ_pnoy_/θπ_horse_), ROH_pony_, θπ, and gene models. Genes associated with body development are highlighted in blue.

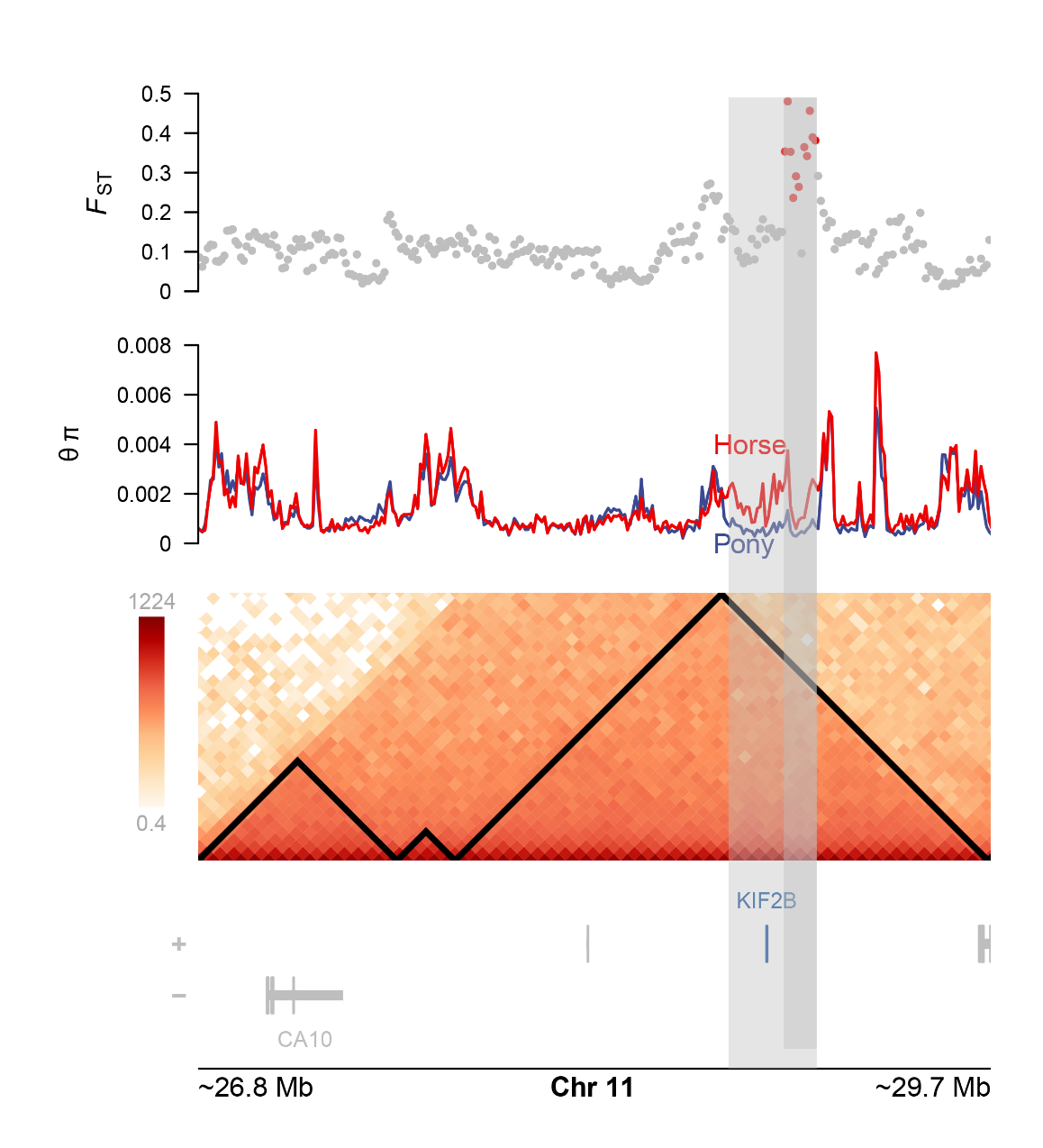

Fig. S20. Candidate selective region on chromosome 11. From top to bottom are *F*_ST_, θπ, Hi-C matrix, and gene models. TADs are indicated with black triangles in the Hi-C matrix.

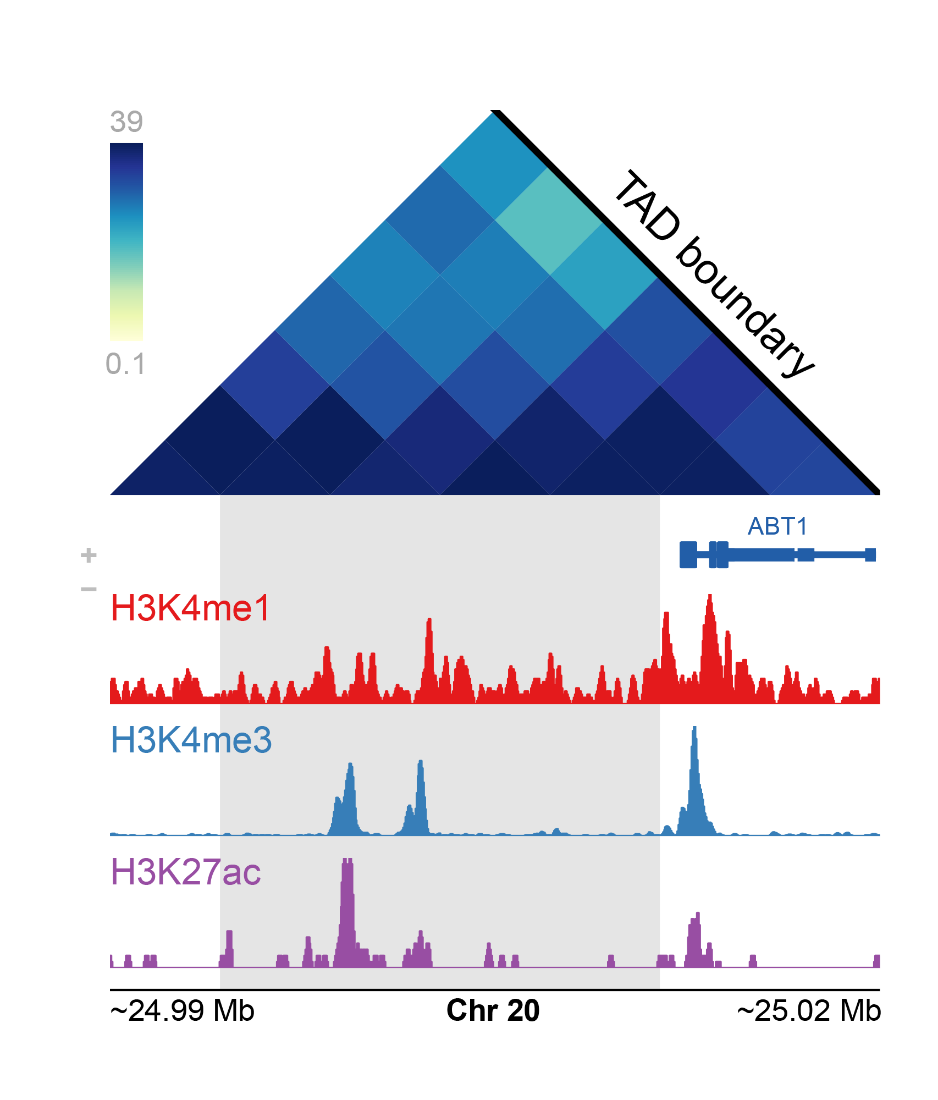

Fig. S21. Candidate selective region on chromosome 20. From top to bottom are the Hi-C matrix, gene models, and histone ChIP-Seq (diaphysis of metacarpal bone) signals. The TAD boundary is marked with a black line in the Hi-C matrix.

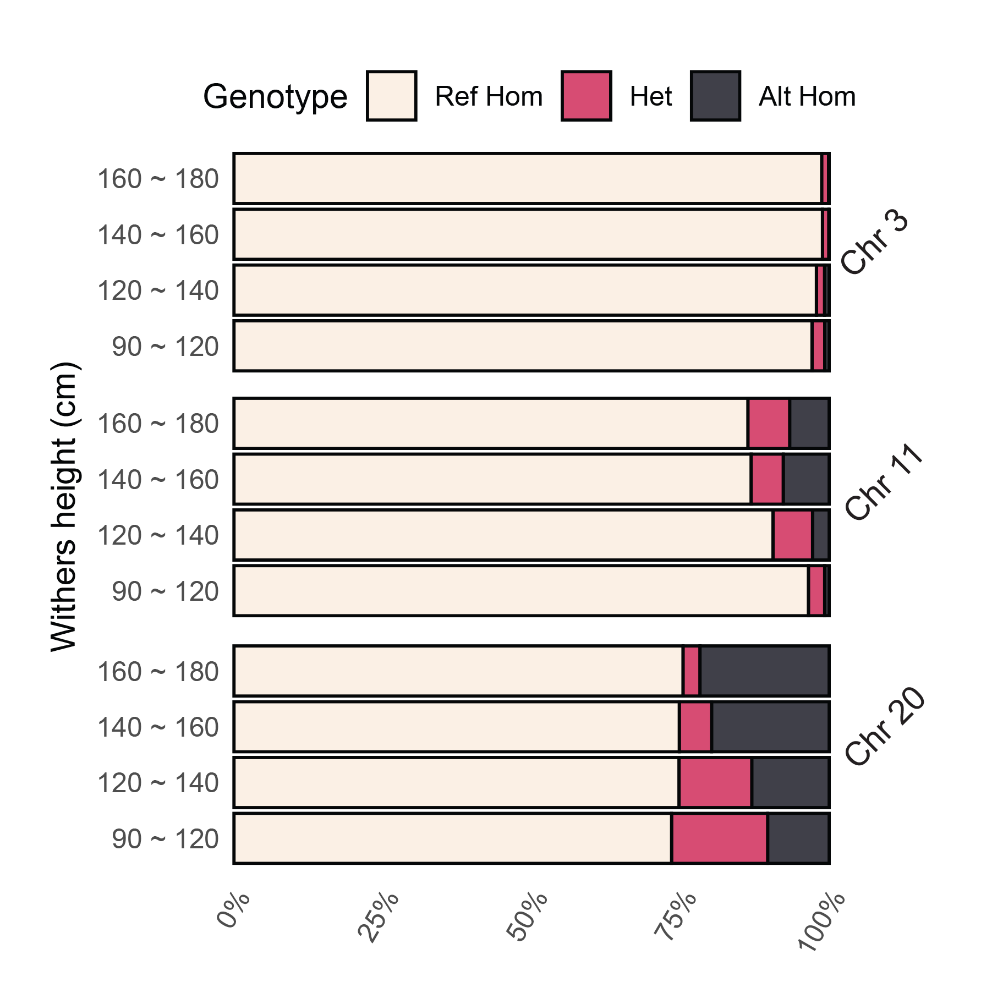

Fig. S22. Genotype frequency of candidate regions at different withers heights located on chromosomes 3, 11, and 20.

Table S1. GenomeScope profiling (31-mer) of the EquCab3.0 and DeBao1.0 assemblies.

|  | **EquCab3.0** | **DeBao1.0** |
| --- | --- | --- |
| Homozygous | 99.81% | 99.71% |
| Heterozygous | 0.19% | 0.29% |
| Genome Haploid Length (bp) | 2,501,801,051 | 2,442,151,487 |
| Genome Repeat Length (bp) | 299,099,790 | 277,828,025 |
| Genome Unique Length (bp) | 2,202,701,261 | 2,164,323,463 |
| Model Fit (bp) | 99.07% | 98.94% |
| Read Error Rate | 0.16% | 0.19% |

Table S2. Assembly statistics throughout the genome profiling stage of the DeBao1.0 assembly and the final EquCab3.0 assembly. The contig N50 was calculated by breaking the genome into contigs at the gap region.

Please refer to details in the file labeled as [**Supplementary Table 2.xlsx**].

Table S3. Summary of repetitive elements, including interspersed repeats and low complexity DNA sequences, identified using RepeatMasker.

|  | **Number of**  **elements** | **Length occupied**  **(bp)** | **Percentage of**  **Sequence (%)** |
| --- | --- | --- | --- |
| Retroelements | 2,224,151 | 800,088,192 | 32.81 |
| SINEs: | 470,777 | 72,714,244 | 2.98 |
| LINEs: | 1,108,026 | 534,035,162 | 21.90 |
| LTR elements: | 645,348 | 193,338,786 | 7.93 |
| DNA transposons | 333,182 | 77,004,990 | 3.16 |
| Unclassified | 656,334 | 101,965,634 | 4.18 |
| Rolling-circles | 1,822 | 479,160 | 0.02 |
| Small RNA | 143,368 | 26,258,168 | 1.08 |
| Satellites | 8,120 | 5,699,367 | 0.23 |
| Simple repeats | 372,965 | 16,795,131 | 0.69 |
| Low complexity | 73,187 | 3,819,876 | 0.16 |

Table S4. Sample information in this study. Samples sequenced in this study are indicated in the column of BioProject. And samples used in each analysis section are also indicated.

Please refer to details in the file labeled as [**Supplementary Table 4.xlsx**].

Table S5. Information of approximate withers height and origin for each horse breed.

| **Breed** | **Origin** | **Withers height (cm)** |
| --- | --- | --- |
| Connemara pony | Ireland | 128 to 148 |
| Arabian horse | Arabian peninsula | 145 to 155 |
| Sorraia | Portugal and Germany | 144 to 148 |
| Hanoverian horse | Germany | 160 to 175 |
| Dülmener pony | Germany | 122 to 132 |
| Standardbred | United States | 160 to 165 |
| Thoroughbred | England | 157 to 173 |
| Shetland pony | Shetland Islands, Scotland | 87 to 107 |
| Akhal-Teke | Turkmenistan | 150 to 160 |
| Andalusian horse | Spain, Iberian Peninsula | 150 to 169 |
| Criollo horse | Pampas (Argentina, Uruguay to Brazil) | 140 to 150 |
| Debao pony | China | 90 to 103 |
| Friesian horse | Netherlands | 152 to 173 |
| Mongolian horse | Mongolia | 130 to 140 |
| Przewalski's horse | Mongolia | 122 to 142 |
| Jeju horse | South Korea | 110 to 130 |
| Saxon-Thuringian heavy warmblood | Saxony to Thuringia | 157 to 165 |
| Italian Trotter | Italy | 145 to 160 |
| Lipizzan | Austria, Croatia, Hungary, to Slovenia | 155 to 165 |
| American Quarter Horse | United States | 145 to 165 |
| Shire horse | United Kingdom | 163 to 183 |
| Lusitano | Portugal | 155 to 165 |
| Warmblood | NA | 160 to 170 |
| Tibetan pony | Tibet | 122 to 127 |
| Yakutian horse | Yakutia (Russia) | 135 to 141 |
| Freiberger | Switzerland | 150 to 160 |
| Friesian dwarf | Netherlands | 112 to 120 |
| German Riding Pony | Germany | 138 to 148 |
| Welsh Pony | Wales | 122 to 137 |
| Morgan horse | United States | 145 to 157 |
| Holsteiner | Schleswig-Holstein, Germany | 163 to 173 |
| Oldenburger | Germany | 165 to 175 |
| Haflinger | Austria, Italy | 138 to 150 |
| Swiss Warmblood | Switzerland | 155 to 168 |
| Dutch Warmblood | Netherlands | 160 to 175 |
| Miniature horse | United States | 86 to 97 |
| Percheron | France | 160 to 185 |
| Tennessee Walking Horse | Tennessee, USA | 150 to 173 |
| Chakouyi horse | China | 127 to 132 |
| Mangalarga Marchador | Brazil | 146 to 152 |
| Icelandic horse | Iceland | 132 to 142 |
| Noriker | Austria | 155 to 165 |
| Trakehner | Prussia | 160 to 170 |
| American Paint Horse | United States | 148 to 160 |
| Traber | United States, France, Germany, England | 150 to 160 |
| Württemberger | Germany | 160 to 175 |
| Westphalian horse | Westphalia, Germany | 160 to 172 |
| Bavarian Warmblood | Germany (south) | 162 to 170 |
| Erlunchun horse | China | 122 to 130 |
| Baise horse | China | 112 to 117 |

Table S6. Sequencing and alignment statistics of WGS samples used in this study.

Please refer to details in the file labeled as [**Supplementary Table 6.xlsx**].

Table S7. Details of the variant rate across the chromosomes.

| **Chromosome** | **Length**  **(bp)** | **Variants** | **Variants rate**  **(bp/variant)** |
| --- | --- | --- | --- |
| 1 | 186,543,716 | 2,594,675 | 71 |
| 2 | 120,552,855 | 1,669,909 | 72 |
| 3 | 119,789,106 | 1,633,222 | 73 |
| 4 | 109,207,483 | 1,599,204 | 68 |
| 5 | 96,479,276 | 1,301,314 | 74 |
| 6 | 87,288,368 | 1,226,430 | 71 |
| 7 | 100,377,767 | 1,375,986 | 72 |
| 8 | 95,742,220 | 1,483,288 | 64 |
| 9 | 84,760,415 | 1,151,371 | 73 |
| 10 | 84,864,005 | 1,221,516 | 69 |
| 11 | 61,402,275 | 801,957 | 76 |
| 12 | 35,324,475 | 921,869 | 38 |
| 13 | 43,811,441 | 695,347 | 63 |
| 14 | 93,526,107 | 1,260,978 | 74 |
| 15 | 92,014,976 | 1,286,833 | 71 |
| 16 | 88,975,660 | 1,160,555 | 76 |
| 17 | 81,184,063 | 1,141,500 | 71 |
| 18 | 82,554,787 | 1,173,488 | 70 |
| 19 | 62,973,423 | 909,846 | 69 |
| 20 | 64,775,031 | 1,347,985 | 48 |
| 21 | 58,446,468 | 887,585 | 65 |
| 22 | 50,341,792 | 697,249 | 72 |
| 23 | 55,512,843 | 809,575 | 68 |
| 24 | 47,484,312 | 690,780 | 68 |
| 25 | 39,825,224 | 574,466 | 69 |
| 26 | 42,089,453 | 668,467 | 62 |
| 27 | 40,797,346 | 689,609 | 59 |
| 28 | 46,908,431 | 686,662 | 68 |
| 29 | 34,839,497 | 555,537 | 62 |
| 30 | 30,474,219 | 478,557 | 63 |
| 31 | 26,152,973 | 383,735 | 68 |
| X | 127,501,132 | 1,387,770 | 91 |
| Y | 8,013,960 | 24,757 | 323 |
| mt | 16,616 | 59 | 281 |

Table S8. Variant distribution across different genomic regions.

| **Type** | **Count** | **Percent**  **(%)** |
| --- | --- | --- |
| Downstream | 1,938,042 | 4.46 |
| Exon | 728,123 | 1.68 |
| Intergenic | 26,318,366 | 60.60 |
| Intron | 11,882,491 | 27.36 |
| Acceptor (Splice site) | 2,194 | 0.01 |
| Donor (Splice site) | 2,653 | 0.01 |
| Region (Splice site) | 42,232 | 0.10 |
| Transcript | 9 | 0 |
| Upstream | 1,912,732 | 4.40 |
| 3′ UTR | 481,659 | 1.11 |
| 5′ UTR | 123,572 | 0.29 |

Table S9. Top 20 principal components in the horse population’s principal component analysis (PCA).

Please refer to details in the file labeled as [**Supplementary Table 9.xlsx**].

Table S10. Admixture components of the horse population for *K* values ranging from 2 to 20.

Please refer to details in the file labeled as [**Supplementary Table 10.xlsx**].

Table S11. Read coverage and sequencing depth on chromosomes 1, X, and Y.

Please refer to details in the file labeled as [**Supplementary Table 11.xlsx**].

Table S12. Y chromosome haplotypes (HTs) of 177 male horse individuals.

Please refer to details in the file labeled as [**Supplementary Table 12.xlsx**].

Table S13. PHATE derived from the top 98 principal components of the PCA.

Please refer to details in the file labeled as [**Supplementary Table 13.xlsx**].

Table S14. Migration statistics with the optimal three migration events. The top 2 mixture events were show in Fig. 4D.

| **Pair** | **MeanW** | **MeanWj** | **MeanSEj** | **Maxpval** | **N** |
| --- | --- | --- | --- | --- | --- |
| Yakutian horse->Shetland pony | 0.236643 | 0.238659 | 0 | 0.00E+00 | 30 |
| Yakutian horse->Debao pony | 0.144504 | 0.144121 | 0.009776 | 0.00E+00 | 30 |
| Yakutian horse->Przewalski's horse | 0.121253 | 0.120915 | 0.008637 | 0.00E+00 | 30 |

Table S15. Selective regions across autosomes with the top 1% *F*_ST_ values.

Please refer to details in the file labeled as [**Supplementary Table 15.xlsx**].

Table S16. Selective regions across autosomes with the top and bottom 1% π ratio (θπ_pnoy_/θπ_horse_).

Please refer to details in the file labeled as [**Supplementary Table 16.xlsx**].

Table S17. Selective regions across autosomes with the top and bottom 1% XP-EHH values (pony vs. horse).

Please refer to details in the file labeled as [**Supplementary Table 17.xlsx**].

Table S18. KEGG pathway enrichment analysis of genes within the selective regions identified by *F*_ST_, π ratio (θπ_pnoy_/θπ_horse_), and XP-EHH (pony vs. horse).

Please refer to details in the file labeled as [**Supplementary Table 18.xlsx**].
